# Hierarchical lineage architecture of human and avian spinal cord revealed by single-cell genomic barcoding

**DOI:** 10.1101/2025.10.24.684328

**Authors:** Giulia L. M. Boezio, Jasper R. L. Depotter, Thomas J. R. Frith, Arthur Radley, Stephanie Strohbuecker, Ana C. Cunha, Michael Howell, James Briscoe

## Abstract

The formation of neural circuits depends on the precise spatial and temporal organisation of neuronal populations during development. In the vertebrate spinal cord, progenitors are patterned into molecularly defined domains, but how lineage relationships shape neuronal diversity and function has remained unclear. Here, we combine genomic barcoding with single-cell RNA sequencing in chick and human embryos to generate cell-type-resolved clonal maps. We find that spinal neurogenesis follows a hierarchical organisation in which the neural tube first partitions into five broad subdivisions that resolve into the eleven progenitor domains generating the cardinal neuronal classes. This bifurcating architecture implies a patterning mechanism of sequential binary decisions. The most prominent lineage restriction occurs at the embryonic alar-basal boundary, separating sensory-processing from motor-control circuits. Individual progenitors generate neurons across multiple temporal waves while remaining constrained within their lineage subdivision, demonstrating persistence of spatial identity despite temporal competence changes. Among sensory populations, we identify two developmental routes, via unifated or bifated progenitors, to pain- and itch-processing interneurons. These principles are conserved between chick and human, with clonal analysis in human embryos revealing that most fate choices are resolved by six weeks post-conception. Together, these findings provide a framework for spinal cord development and reveal lineage compartmentalisation as a fundamental principle in neural circuit assembly and evolution.

## Introduction

The spinal cord transforms sensation into action. The processing of sensory inputs and coordination of motor output rely on a remarkable diversity of neuronal types organised in precisely wired neural circuits. This functional complexity is initiated during embryonic development with the spatially and temporally organised specification of neural progenitor cells that generate the post-mitotic neuronal subtypes involved in spinal circuits (*1*). Recent single-cell transcriptomic studies have documented extensive molecular heterogeneity within neuronal and glial populations and revealed that both spatial and temporal mechanisms contribute to functional specialisation (*2–6*). Yet these studies do not reveal cell lineage directly, raising fundamental questions about the cellular organisation underpinning this molecular diversification: What are the lineage relationships between developing cell types? Do progenitor domains organise into broader lineage pedigrees, or are molecularly-distinct domains lineage-restricted? Do transcriptionally similar neurons share the same origin? How does developmental timing shape individual progenitor competence? Does developmental compartmentalisation mirror functional circuit organisation? Addressing these questions is essential for understanding how patterning shapes circuit assembly and has broad implications for evolution, development, disease, and regenerative medicine.

These questions arise against a backdrop of well-established principles describing how progenitor and neuronal subtype identity are specified along the dorsoventral axis of the spinal cord. Signalling molecules, including Sonic Hedgehog (SHH) and Bone Morphogenetic Proteins (BMP), emanating from opposing sources establish transcriptional programmes that define eleven cardinal progenitor domains (Fig. 1A) through the combinatorial expression of a suite of transcription factors (TFs) (*1*, *7*) (Fig. 1A). Loss-of-function and gain-of-function experiments have highlighted the necessity of key transcription factors for fate specification, including Msx1, Atoh1, Ptf1a, Pax7, Pax6, Dbx1/2, Olig2, and Nkx2.2 (*8–21*). The action of these proteins determines the neurons produced: motor neurons (MNs) and locomotor interneurons (V0-V3) from ventral domains, sensory neuron subtypes (dI1-dI6) from dorsal regions, and different types of glial cells from distinct domains at later developmental stages (*22–26*). Temporal patterning adds another layer of complexity (*26–28*). Spinal cord neurogenesis proceeds through sequential waves marked by distinct TF combinations that specify early-born versus late-born neurons (*2*, *29*). This temporal code amplifies neuronal diversity by creating subtypes within each cardinal neuronal class based on time of birth (*28*). Yet, despite this detailed molecular understanding, how individual progenitor cells contribute to neuronal and glial diversity, and ultimately to functional circuit assembly, remains largely unresolved.

**Figure 1.**
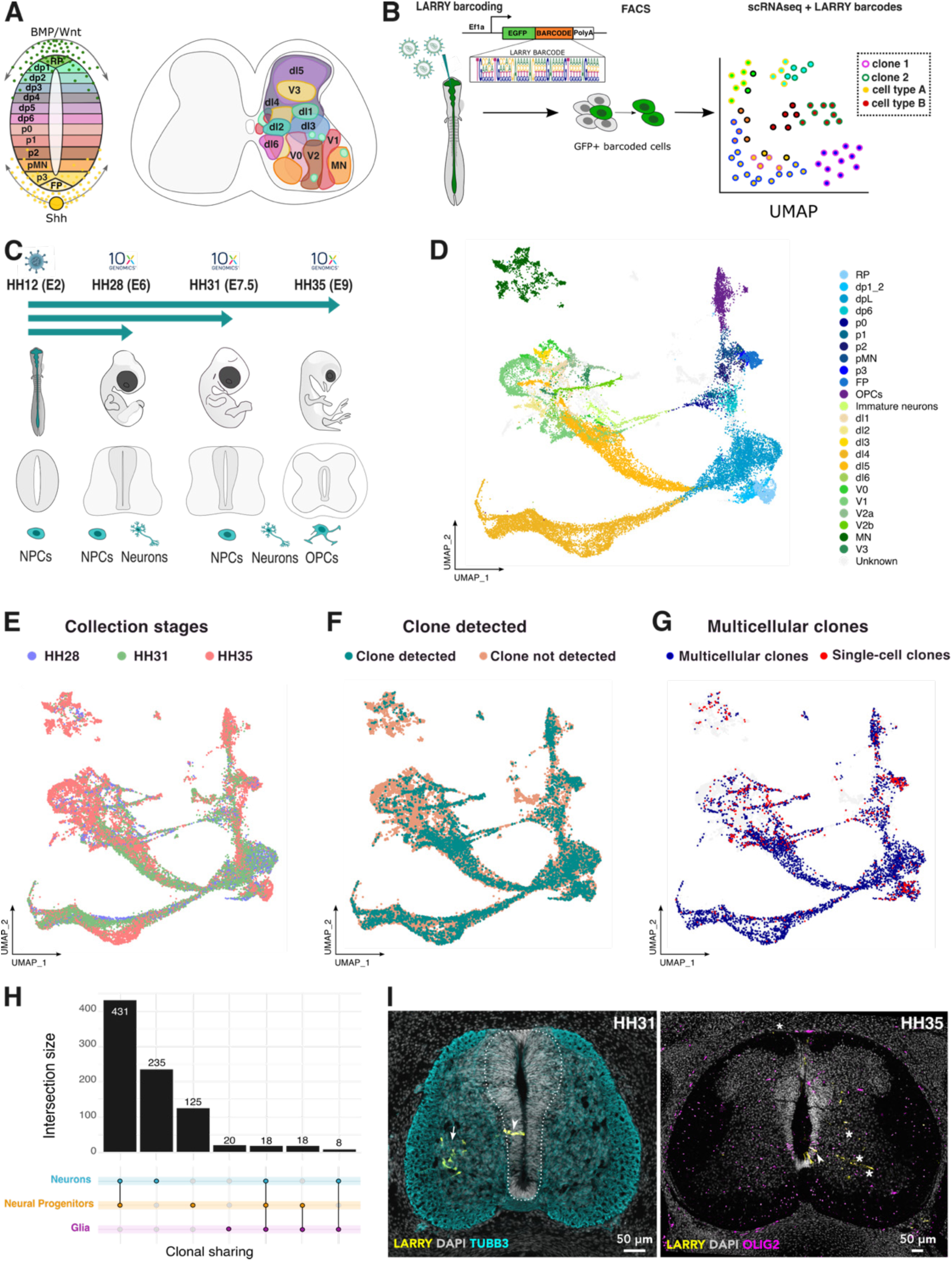
Lineage tracing of spinal cord development using LARRY barcoding A. Schematic representation of dorsal-ventral (DV) progenitor domains in transverse sections of the neural tube (left) and corresponding neuronal populations in the mature spinal cord (right). B. Overview of the LARRY barcoding workflow for lineage tracing. C. Experimental timeline of LARRY barcoding, showing viral infection and sample collection stages (and neural tube cell type composition). D. UMAP of integrated single-cell transcriptomes, coloured by assigned cell types. E–G. UMAPs coloured by collection developmental stage (E), barcode-expressing cells (F), and cells in multicellular clones (G). H. UpSet plot displaying co-occurrence of cell types within individual clones. Top: bar plot quantifying clone combinations; bottom: graphical table showing the specific cell type combinations. I. Confocal images of transverse sections of chick neural tube stained for EGFP (yellow; barcoded cells), TUBB3 (cyan; neurons), and OLIG2 (magenta; pMN progenitors and OPCs), highlighting barcoded progenitors (arrowheads), neurons (arrows), and oligodendrocytes (asterisks) at two collection developmental stages (HH31; HH35).

Previous lineage tracing studies using Cre-lines, viral vectors, and fluorescent dyes established the conceptual framework for understanding progenitor dynamics (*30–39*), but were limited by low throughput, restricted molecular annotation, and uncertainty over true clonality. Key questions remain unresolved: for example, whether two cell types arise from the same individual progenitor (e.g. motor neurons and oligodendrocytes), or when in development progenitors commit to specific neuronal classes.

Recent advances in single-cell technologies offer new possibilities for unbiased, high-resolution lineage tracing in complex tissues (*40–42*). Genomic barcoding techniques, such as LARRY (<u>L</u>ineage <u>A</u>nd <u>R</u>NA <u>R</u>ecover<u>Y</u>), enable simultaneous tracking of thousands of progenitor lineages while preserving molecular identity through single-cell RNA sequencing (*43*). These methods overcome the limitations of traditional approaches by providing comprehensive, molecularly annotated lineage maps at unprecedented scale and resolution. In addition, because they do not require the generation of complex transgenic lines, they hold the potential to be applied broadly across species, including to human tissue. While early *in vivo* applications have demonstrated the feasibility of these approaches in mouse models (*44–46*), their use in complex tissues has remained limited, and resolving fine-grained lineage relationships in highly heterogeneous systems has remained a challenge.

Here, we adapt high-throughput genomic barcoding for comprehensive lineage analysis in both chick and human spinal cord development *in vivo*. Our analysis reveals an unexpected modular hierarchical organisation, in which the neuroepithelium is initially partitioned into five discrete *lineage subdivisions*, which lie hierarchically above the canonical 11 progenitor domains. The subdivisions align with broad neural circuit function, with the major dorsal-ventral lineage restriction corresponding to the embryonic alar-basal boundary that separates sensory and motor systems. Furthermore, we reveal how dual developmental routes generate functional diversity within sensory interneuron populations. Critically, these lineage principles are conserved between chick and human. These results offer a framework for understanding how early embryonic patterning constrains neural circuit assembly and provide insight into the evolutionary origin of neural tube organisation. By revealing how diversity is encoded in the lineage of spinal neurons, we provide a conceptual framework that may extend to other regions of the nervous system and illuminate conserved principles across species.

## Results

### High-resolution *in vivo* lineage reconstruction in the chick spinal cord

To map spinal neuron lineages *in vivo* (Fig. 1A), we adapted the genomic barcoding technique LARRY (*43*) to chick (*Gallus gallus*) embryos (Fig. 1B). We injected a high-diversity lentiviral barcode library (Material & Methods) into the neural tube lumen at Hamburger-Hamilton (HH) stage 12 (embryonic day 2, comparable to mouse E8-8.5 (*47*) and human Carnegie Stage (CS)10-11 (https://hdbratlas.org/comparison-HvM.html)) (Fig. 1C). At this stage, the closed neural tube primarily comprises neural progenitors with neurogenesis in the ventral domains just starting (Fig. 1C, S1A). Transcription factors patterning neural progenitors begin expression at these stages, initiating the process that establishes the eleven canonical dorsoventral progenitor domains ((*16*, *22*); Fig. S1A).

We collected barcoded cells from the embryos’ brachial region at three developmental timepoints: HH28, HH31-32, and HH35 (equivalent to mouse E12-E14) (Fig. 1C, E). This sampling window captures the transition from active neurogenesis to early gliogenesis (Fig. S1B-C). We processed 17 barcoded samples (n=4, 7, and 6 per timepoint, respectively) alongside unlabelled controls for single-cell RNA sequencing. Cell type annotation identified all expected cardinal neuron classes, including V3, motor neurons (MNs), V2a/b, V1-0, and dorsal interneurons (dI)1-6, plus their cognate progenitors transitioning to ependymal fates and stage-specific oligodendrocyte precursor cells (OPCs) (Fig. 1D, S1B-H). Astrocytes were not detected, though sparse *GFAP* expression appeared at HH35, consistent with previous reports ((*48*, *49*), Fig. S1H).

Using a newly established computational pipeline (Materials and Methods), we identified approximately 1,500 clones, including 867 multicellular clones, representing 84% of labelled cells (Fig. 1F-G, Fig S2A). Clone size increased over time (from 2.7 to 4.5 cells per clone, Fig. S2B), consistent with continuing cell division in the progenitor populations. Quantification of intact clones in tissue sections revealed average clone sizes of 7 cells in ventral and 15 in dorsal regions (Fig. S2C-E), reflecting the earlier onset and shorter duration of ventral neurogenesis and the longer progenitor expansion phase of dorsal domains (*47*). Comparing these *in situ* measurements to clone sizes recovered from the LARRY transcriptomic dataset suggests that scRNA-seq captures approximately 20% of the cells in each clone (Fig. S2C-E). Clones distributed broadly across dorsoventral and mediolateral axes, frequently linking neural progenitors with their differentiated progeny, confirming the robustness of our lineage reconstruction approach (Fig. 1H-I, S2F-G).

### The neural tube comprises five lineage subdivisions

We first investigated how lineage relationships are spatially organised within the developing spinal cord. We quantified how often distinct cell types shared barcodes, providing a readout of common developmental origin. By projecting these relationships along the dorsoventral axis, we constructed a global map of lineage architecture (Fig. 2A). To account for population size differences, we calculated clonal coupling z-scores, which quantify the extent of barcode sharing between two cell types compared to a random expectation (Fig. S3A-B) (*44*, *46*, *50*). Positive values indicate more frequent co-occurrence in clones than expected by chance, while negative values indicate lower-than-expected clone sharing. To identify broader patterns of lineage organisation, we then calculated clonal coupling correlation scores to identify groups of cell types that are clonally related (Fig. 2A; (*46*, *50*); Materials and Methods). In these matrices, diagonal values indicate how often cells of the same type co-occur in clones. High values reflect strong within-type clonal pedigrees, while lower values suggest asymmetric divisions or temporal diversification. Off-diagonal high values highlight shared clonal origins between distinct cell types, reflecting common lineage ancestry.

**Figure 2.**
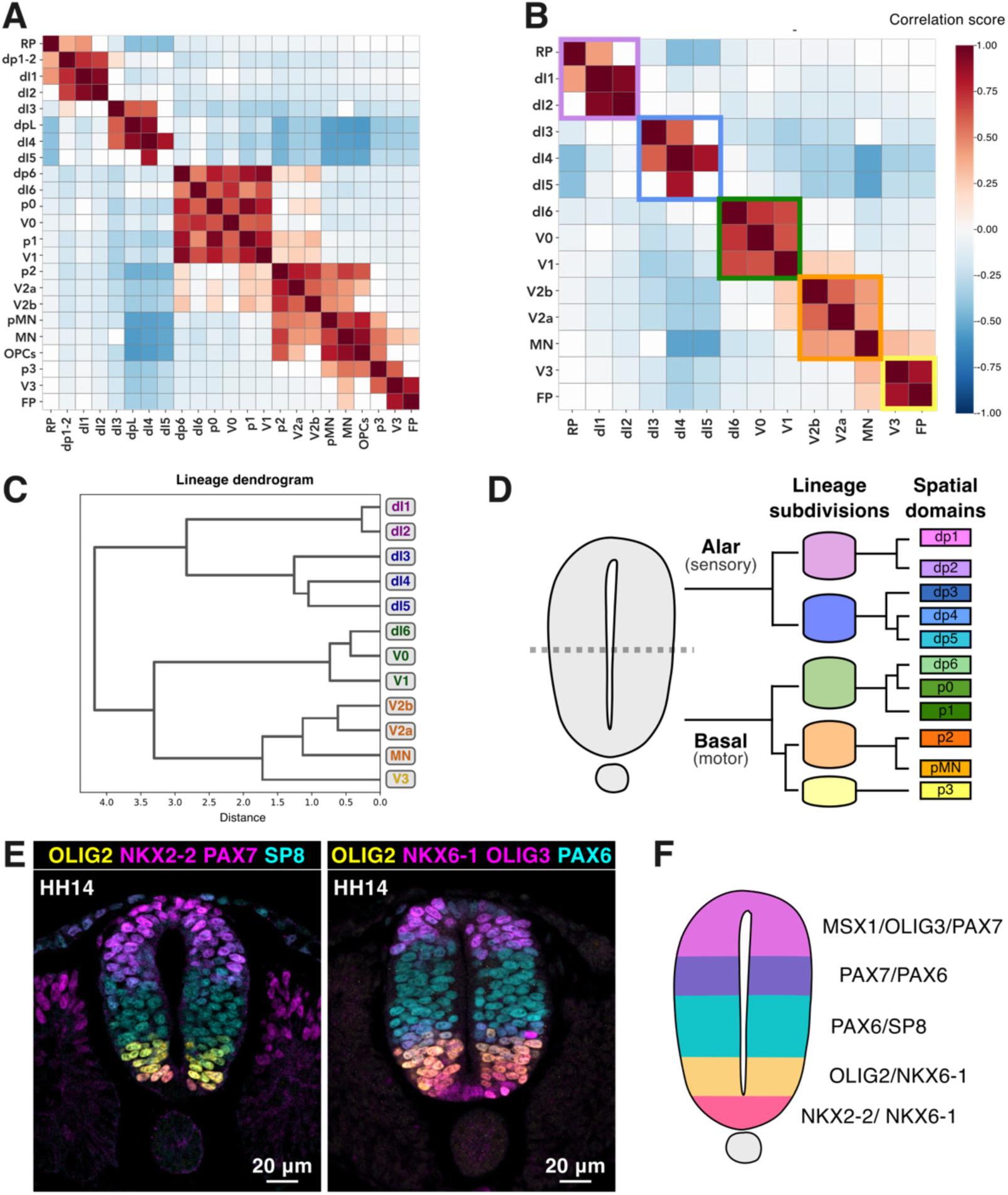
**The neural tube comprises five lineage subdivisions aligning with functional circuit organization** A–B. Heatmaps of clonal coupling correlation scores for all cell types (A) and for neurons only (B), with five lineage subdivisions highlighted. Diagonal values indicate how frequently cells of the same type are found in the same clone; off-diagonal blocks highlight shared progenitor relationships between different cell types. C. Dendrogram showing hierarchical clustering of clonal coupling correlation scores for all neurons, revealing the nested structure of lineage relationships. D. Schematic illustrating the progressive refinement of lineage potential during spinal cord development. Early progenitors partition into lineage subdivisions, which give rise to spatially defined progenitor domains, each producing distinct neuronal classes. The lineage architecture mirrors the classic alar–basal division separating dorsal (sensory) and ventral (motor) territories (left). E-F. Confocal images and diagram of HH14 transverse sections stained for transcription factors defining each compartment: OLIG2, NKX2-2, PAX7, SP8, NKX6-1/2, OLIG3, and PAX6.

Most frequently, barcodes are shared between cells of the same progenitor domain and neuronal subtype (Fig. 2A, diagonal line). This pattern suggests that spatial fate bias exists early in development and that positional identity persists throughout neurogenesis. In addition, the data confirmed several longstanding developmental models of neuronal specification. For example, V2a and V2b interneurons, previously shown to share a common domain origin (*38*, *51*, *52*), consistently co-occurred within individual clones, supporting the existence of a bifated V2 progenitor (Fig. 2A, S3A).

Previous retroviral tracing studies identified clonal lineages that spanned cells in spatially adjacent regions (*37*). We also observed neighbouring cell types sharing lineages, but the higher resolution and molecular annotation of our approach uncovered an unexpected structure within these domain- spanning relationships. Rather than clones being equally likely to comprise any pair of neighbouring cell types, clonal relationships clustered into discrete, spatially coherent blocks. We refer to these as *lineage subdivisions* (Fig. 2B). These subdivisions group cells that are strongly clonally coupled, documenting a previously unappreciated higher-order organisation of progenitor output, a lineage-based scaffold that sits above the canonical progenitor domains.

Five distinct subdivisions were apparent from dorsal to ventral: roof plate and dI1–2 neurons (with their progenitor domains); followed by dI3–5; then dI6 to V1 (spanning dp6–p1); a subdivision encompassing p2, pMN (including V2a/b interneurons, motor neurons, and OPCs); and finally, floor plate and V3 neurons. The p2/V2 domain and motor neurons showed broader clonal overlap with neighbouring subdivisions, suggesting a transitional identity (Fig. 2A-B).

Hierarchical clustering suggests that these subdivisions are structured through sequential bifurcations (Fig. 2C, S3C). The nested arrangement of lineage relationships is consistent with a model of progressive lineage restriction, with the most prominent bifurcation separating dorsal and ventral progenitors at the dI5-dI6 boundary (Fig. 2D). Within the dorsal compartment, dI1-2 and dI3-5 arise from distinct lineage branches. In the ventral spinal cord, progenitors subdivide into dp6-p1 and then p2-pMN and p3-floor plate (FP) groups. This organisation is consistent with recent chromatin accessibility profiling showing distinct regulatory landscapes between p3/FP and p2/pMN progenitors (*53*), and with dynamical modelling suggesting that progenitor diversification occurs through discrete bifurcations rather than by sequential induction at a series of monotonic morphogen thresholds (*54*).

To investigate whether the lineage architecture corresponds to early molecular differences, we examined transcription factor expression a few hours after barcoding (HH14). Each lineage subdivision was associated with a distinct combination of transcription factors, including MSX1, OLIG3, PAX7, PAX6, SP8, NKX6-1/2, OLIG2, and NKX2-2, which pattern the 11 progenitor domains at later stages (Fig. 2E; Fig. S3D). Notably, at this earlier stage, these factors were expressed in broader patterns that overlapped with the five subdivisions (Fig. 2F), suggesting that the molecular priming of lineage potential precedes the emergence of discrete progenitor domains and the onset of neurogenesis.

### Hierarchical lineage subdivisions align with functional circuit organization

The hierarchical organisation of lineage subdivisions raised the question of whether this developmental architecture has functional relevance. To explore this, we asked whether subdivisions align with known aspects of circuit architecture. The hierarchical clustering suggested that the most prominent boundary occurs between dI5 and dI6 (Fig. 2C, S3C), corresponding to the classic alar–basal division that separates dorsal (sensory) from ventral (motor) territories. To investigate this boundary, we focused on so-called class B interneurons ((*55*); dI4-dI6), which span the critical transition zone. These neurons are highly abundant and play key roles in sensory processing and motor control, yet their developmental origins and regulatory mechanisms remain incompletely understood (*5*, *6*, *21*, *56–58*).

dI6 neurons are usually classified as dorsal, based on their nomenclature, gene expression, and derivation *in vitro* using protocols that generate dorsal neuronal subtypes (*6*, *21*, *59*, *60*). However, our clonal analysis showed that dI6 neurons frequently co-occur in the same clones as ventral V0 and V1 interneurons and rarely share lineages with dorsal dI4s or dI5s (Fig. 2A-B). This clonal relationship mirrors circuit logic: dI6, V0, and V1 neurons all participate in motor control and innervate motor neurons (Fig. S3E) while dI4s and dI5s primarily process sensory inputs (*61–63*). These findings suggest that, despite their usual dorsal designation, dI6 neurons share a ventral lineage and functional trajectory.

The dI5–dI6 lineage boundary exemplifies the logic of the hierarchical subdivision architecture. This boundary marks both a major developmental bifurcation, the most prominent clonal restriction in the dataset, and a functional divide between primarily sensory-processing (dI4–dI5) and motor-control (dI6, V0, V1) systems. The alignment between lineage organisation and circuit function suggests that early binary progenitor fate decisions establish not only cell type diversity but also the modular functional architecture of spinal circuits (Fig. 2D).

### Temporal diversification of neurons and glia within lineage subdivisions

Having defined the spatial structure of spinal cord lineages, we next examined how progenitors generate neuronal diversity within lineage subdivisions. A bifated p2 progenitor gives rise to both V2a and V2b neurons (Fig. 3A). By contrast, V0v and V0D neurons, which are excitatory and inhibitory interneurons involved in left-right coordination, respectively (*63–65*), never co-occur in clones (Fig. 3B), indicating distinct progenitor origins within the broader p0 domain. Their divergent lineage relationships support this separation, with V0v linked to more ventral (e.g., V1, V2) and V0D to more dorsal neurons (e.g., dI4–dI6) (Fig. S4A). This pattern suggests that the p0 domain comprises two molecularly (*66*, *67*) and lineage distinct progenitor pools.

**Figure 3.**
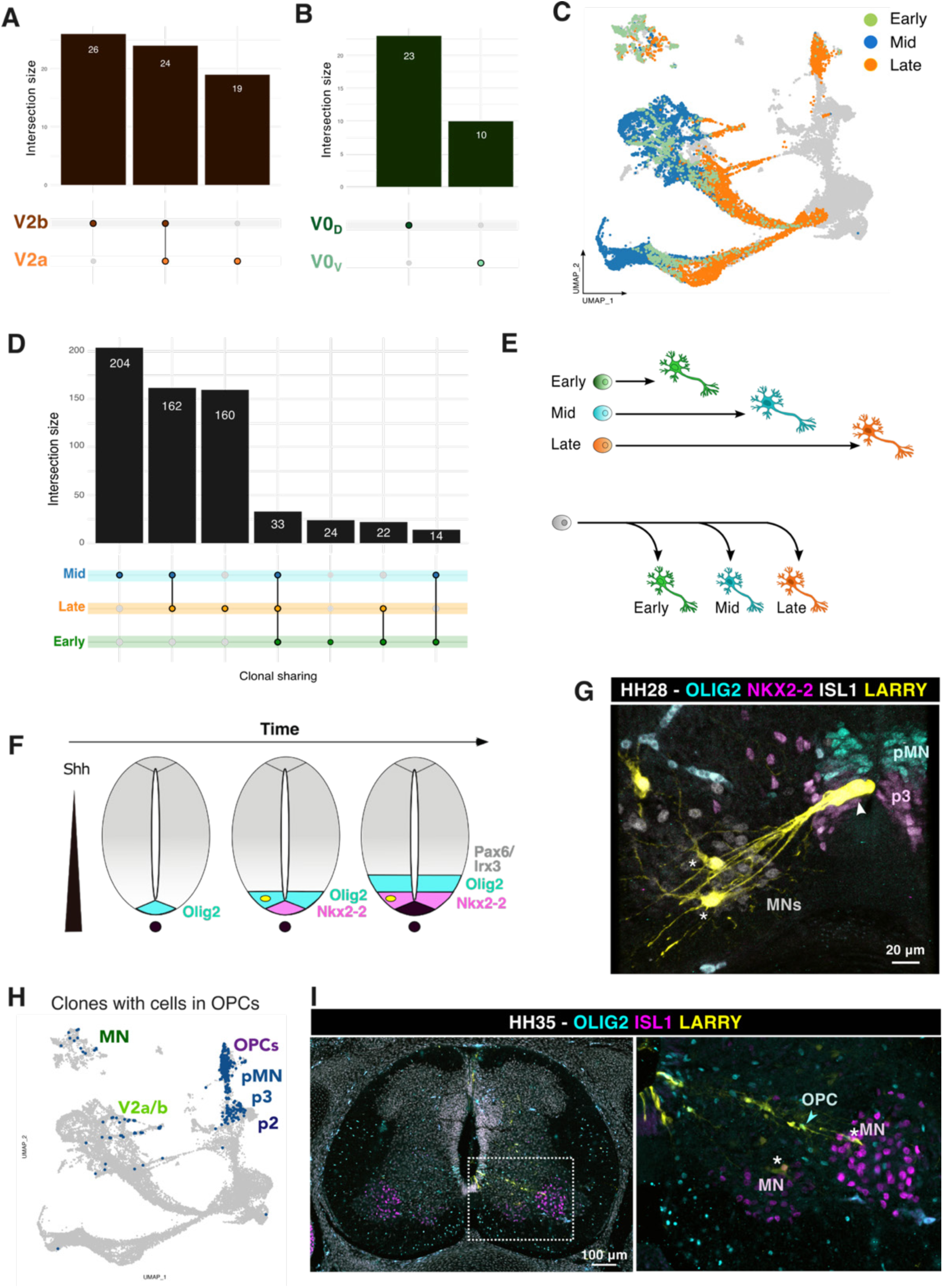
**Temporal diversification of neurons and glia within lineage subdivisions** A-B. UpSet plot displaying co-occurrence of V2a and V2b neurons or lack of co-occurrence of V0V and V0D neurons within individual clones. Top: bar plot quantifying clone combinations; bottom: graphical table showing the specific cell type combinations. C. UMAP coloured by temporal neurogenesis wave D. UpSet plot displaying co-occurrence of cells from different neurogenesis waves within individual clones. Top: bar plot quantifying clone combinations; bottom: graphical table showing the specific cell type combinations. E. Diagram comparing two models of neurogenic timing: distinct progenitor pools vs. multipotent progenitors generating neurons at different times. F. Diagram of progressive ventralisation driven by kinematic transcription factor waves (modified from (*1*)). G. Confocal images of HH28 transverse sections showing one clone containing both MNs (ISL1+, asterisks) and p3 progenitors (NKX2-2+, arrowhead), but no pMN progenitor (OLIG2+). H. UMAP of clones containing at least one OPC, showing clonal relationships among OPCs, pMN, p3, p2, MN, and V2 cells. Colours of cell type names correspond to clusters in Fig. 1D for reference. I. Confocal images of HH35 transverse sections showing clones (EGFP, yellow) comprising MNs (asterisks, ISL1+ magenta) and OPCs (OLIG2+, cyan, arrowheads). MN, motor neuron; OPC, oligodendrocyte progenitor.

The p1 domain displayed a different behaviour. Both Renshaw (*MAFA/B⁺)* and non-Renshaw IaIN-like (*FOXP1/2⁺*) interneurons arise from common individual progenitors (Fig. S4B, C), despite being generated during distinct neurogenic waves (*68–71*). This observation prompted us to test whether the sequential waves of neurogenesis that diversify spinal cord neuronal subtype identity (*28*) occur within single lineages or through separate wave-specific progenitor pools. We classified neurons by temporal birth markers: early (*ONECUT1/2⁺*), mid (*ZFHX3/4⁺*), or late (*NFIA/B⁺, NEUROD2⁺*) born (Fig. 3C, S4D-E), excluding cells with mixed expression. Focusing on mid- and late-wave neurons, which constitute most spinal neurons (*2*), we found that individual progenitors generate neurons from multiple temporal waves (Fig. 3D, S4F-G). This demonstrates that temporal diversification operates within single lineages (Fig. 3E). Critically, multi-wave clones remained largely confined to individual lineage subdivisions (Fig. S4F-G). This constraint suggests that spatial identity persists even as progenitors change their temporal output.

Within clones originating in the ventral neural tube, we observed a tendency towards a ventral shift in their composition over developmental time. For instance, mid-wave neurons were often clonally linked to late-wave neurons emanating from a more ventral domain (Fig. S4H). This bias supports the kinematic wave model, where dynamic transcriptional changes progressively shift progenitor identity toward more ventral fates over time ((*8*); Fig. 3F). Tissue sections from barcoded embryos confirmed this pattern, showing clones containing both MNs, known to arise from the pMN progenitor domain (*14*), and progenitors in the adjacent ventral p3 domain (Fig. 3G). These findings are consistent with a progressive ventralisation of neural progenitors during the neurogenesis window.

These observations prompted us to examine the origins of oligodendrocyte progenitor cells (OPC). OPCs arise in ventral regions of the spinal cord after neurogenesis, suggesting lineage relationships with ventral neurons. We found robust clonal connections between MNs, pMN, and OPCs in both sequencing and imaging data (Fig. 2A, Fig. 3H-I). This supports the long-debated hypothesis (*33*, *35*, *72–79*) that an individual pMN progenitor could directly generate both neuronal and glial lineages through temporal competence switches. Although we cannot exclude the existence of some lineage-restricted cells within the pMN domain (*74*, *78*, *80*), our findings are consistent with the presence of a bifated pMN progenitor pool that switches competence over time, producing first MNs and later OPCs, consistent with retroviral tracing and genetic lineage studies (*35*, *79*). However, OPCs also shared clones with p3 progenitors at the time of collection (Fig. 2A, Fig. 3H). While we cannot comment on the state of a progenitor at the time of gliogenic commitment, this aligns with evidence that both pMN (*Olig2+*) and p3 (*Nkx2.2+*) progenitors contribute to OPC production (*15*, *31*, *81–83*). In addition, clones linking V2a neurons and OPCs were also observed (Fig. 2A, 3H), suggesting that the OPC potential is shared across the subdivision.

These findings reveal a model where progenitor identity shifts progressively toward more ventral fates over time. This temporal plasticity could allow OPCs to emerge from multiple ventral progenitor sources, reconciling previously conflicting lineage models. More broadly, our clonal data show that the temporal progression of neurogenesis can span different neuronal classes while respecting lineage subdivision boundaries.

### Dual developmental routes generate dI4 and dI5 sensory interneurons

dI4s and dI5s comprised the majority of neurons in our dataset (56%; Fig. 4A, B) and formed a cohesive clonal unit despite distinct molecular identities (Fig. 4C, 2B-C). dI4 are inhibitory, while dI5s are excitatory neurons, and both participate in pain, itch, and touch processing circuits (*23*, *61*). Current models propose that dI4s and dI5s arise initially from distinct domains (dp4, Ptf1a+/Ascl1+, and dp5, Ascl1+), before converging later into a shared domain that produces later-born dI4 and dI5 (*55*, *56*, *58*, *84–86*). However, evidence for early separate lineages relies primarily on transcription factor knockout studies.

**Figure 4.**
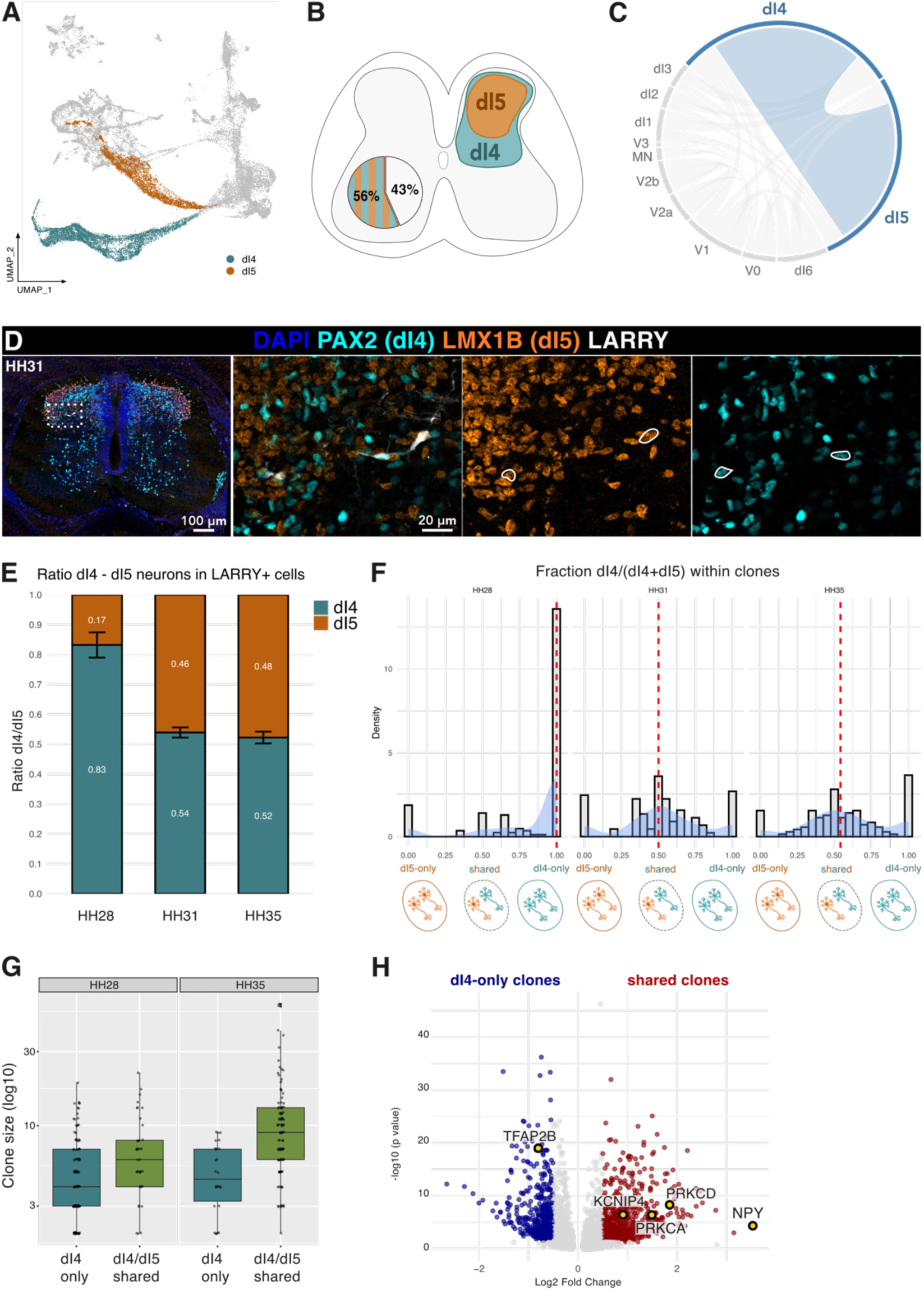
**Lineage origins and trajectories of dorsal sensory neurons** A. UMAP visualisation of dI4 and dI5 neurons. B. Schematics and pie chart showing prevalence of dI4/dI5 neurons among other neurons. C. Circos plot showing the frequency of shared clones between neuronal types. The prominent connection between dI4 and dI5 highlights a strong clonal relationship. D. Confocal images of HH31 transverse spinal cord sections stained for DAPI, PAX2 (dI4), LMX1B (dI5), and EGFP (LARRY+), highlighting shared clones. E. Stacked bar plots of dI4/dI5 neuron ratios across timepoints in barcoded cells. F. Histogram and kernel density estimate (KDE) plot of dI4 / (dI4 + dI5) ratios in clones. Bars represent clone counts within each bin, and the overlaid curve shows a KDE scaled to the histogram for visual comparison. A ratio of 1 indicates clones composed entirely of dI4 neurons, 0 indicates clones composed entirely of dI5 neurons, and 0.5 reflects equal representation. Red dashed line marks median ratio. G. Boxplots of clone size in dI4-only vs. shared clones across timepoints. Shared clones increase in size between HH28 and HH35, while dI4-only clones do not. Volcano plot of differentially expressed genes in dI4 neurons within dI4-only vs. shared clones; T*FAP2B, NPY*, *KCNIP4, PRKCA/D* highlighted.

We found that dI4s and dI5s frequently shared barcodes (Fig. 4C-D, 2B), indicating a shared progenitor origin and challenging the notion of strictly separate early lineages. Temporal analysis revealed dynamic changes in dI4/dI5 ratios (Fig. 4E, S5A), further clarifying these developmental relationships. At HH28, dI4s dominated (80%) while dI5s were underrepresented (20%). By HH35, the ratio equalised to ∼50:50. These dynamic changes in proportions mirror data in developing mouse embryonic spinal cords (Fig. S5B). Individual clone compositions also showed a distinct temporal trend (Fig. 4F). At HH28, clones contained either exclusively dI4 or mixed dI4/dI5 populations, while pure dI5 clones were rare. Over time, the proportion of dI4-only clones decreased, and mixed clones became predominant. In addition, mixed clones grew in size while dI4-only remained stable over time (Fig. 4G).

These findings suggest two possible developmental trajectories (Fig. S5C-D). In the first, a LARRY- labelled progenitor gives rise to two unifated progenitors (Fig. S5C): one producing dI4 (dp4), and another producing dI5 (dp5). A higher rate of differentiation by dp4 cells would explain the early abundance of dI4 and dI4-only clones. The dp5 lineage would exhibit slower or delayed output, resulting in a rise in dI5s at later stages and increased sharing in clones. However, this model would predict a higher prevalence of dI5-only clones at later stages due to the expansion of the dp5 progenitor pool derived from delayed or slower differentiation. An alternative scenario (Fig. S5D), more consistent with the data, implicates two distinct lineages of progenitors: fast-differentiating unifated dp4 progenitors that produce only dI4, and slower-differentiating bifated progenitors that generate both dI4 and dI5. dp4 progenitors differentiate rapidly and are therefore extinguished earlier, explaining the initial dI4 dominance and pure dI4 clones. Shared progenitors differentiate at a slower rate, thereby expanding over time, and contribute most dI4s and dI5s at later stages, explaining the growing mixed clone population (Fig. 4G). We cannot exclude the existence of a distinct dI5-only progenitor population, although it would appear to make only a minor contribution.

This dual-route mechanism raises the possibility that the distinct origins of dI4s correspond to a molecular and functional division within dI4 populations. Consistent with this, dI4s from dI4-only clones expressed higher levels of *TFAP2B* (Fig. 4H), a marker of motor-synergy encoder neurons implicated in reflex coordination (*87*). By contrast, dI4s from mixed clones showed elevated expression of *NPY, PRKCA/D,* and *KCNIP2* (Fig. 4H). Members of the same gene families mark superficial dorsal horn sensory populations in mouse (*5*) implicated in mechanical itch responses and nociceptive reflex plasticity (*88*, *89*). At early stages, nearly all dI4 were *TFAP2B⁺*, whereas *TFAP2B⁻* populations emerged at later stages (Fig. S5E). Conversely, expression of *NPY, PRKCA/D,* and *KCNIP2* was only detected at later stages (Fig. S5F-I). This temporal and clonal segregation suggests that developmental origin shapes functional specialisation within sensory interneurons, opening new avenues to dissect lineage- function relationships in the dorsal spinal cord.

### Human stem cell derived model confirms dI4 and dI5 pedigrees

To test whether the dual-lineage origin of sensory neurons uncovered in chick is conserved in mammals, we turned to a human ESC (embryonic stem cell)–based differentiation model. This offered a tractable system to monitor the emergence of dI4s and dI5s and allowed progenitor dynamics to be tracked at defined time points. This system enabled labelling at different stages, making it possible to compare the pedigrees of early versus late progenitors.

By adapting published protocols (*60*, *90–92*), we generated dorsal PAX7⁺/PAX6⁺/MSX1⁻ progenitors by day 14 (dp3–dp5; Fig. S6A). By day 24, these cultures began producing neurons expressing dI4 and dI5 markers (PAX2 and LMX1B, respectively; Fig. 5A; Fig. S6B), with expression becoming more widespread by day 28 (Fig. 5B). At this stage, additional markers (LHX1/5 for dI4 and TLX3 for dI5) further supported robust subtype specification (Fig. S6C).

**Figure 5.**
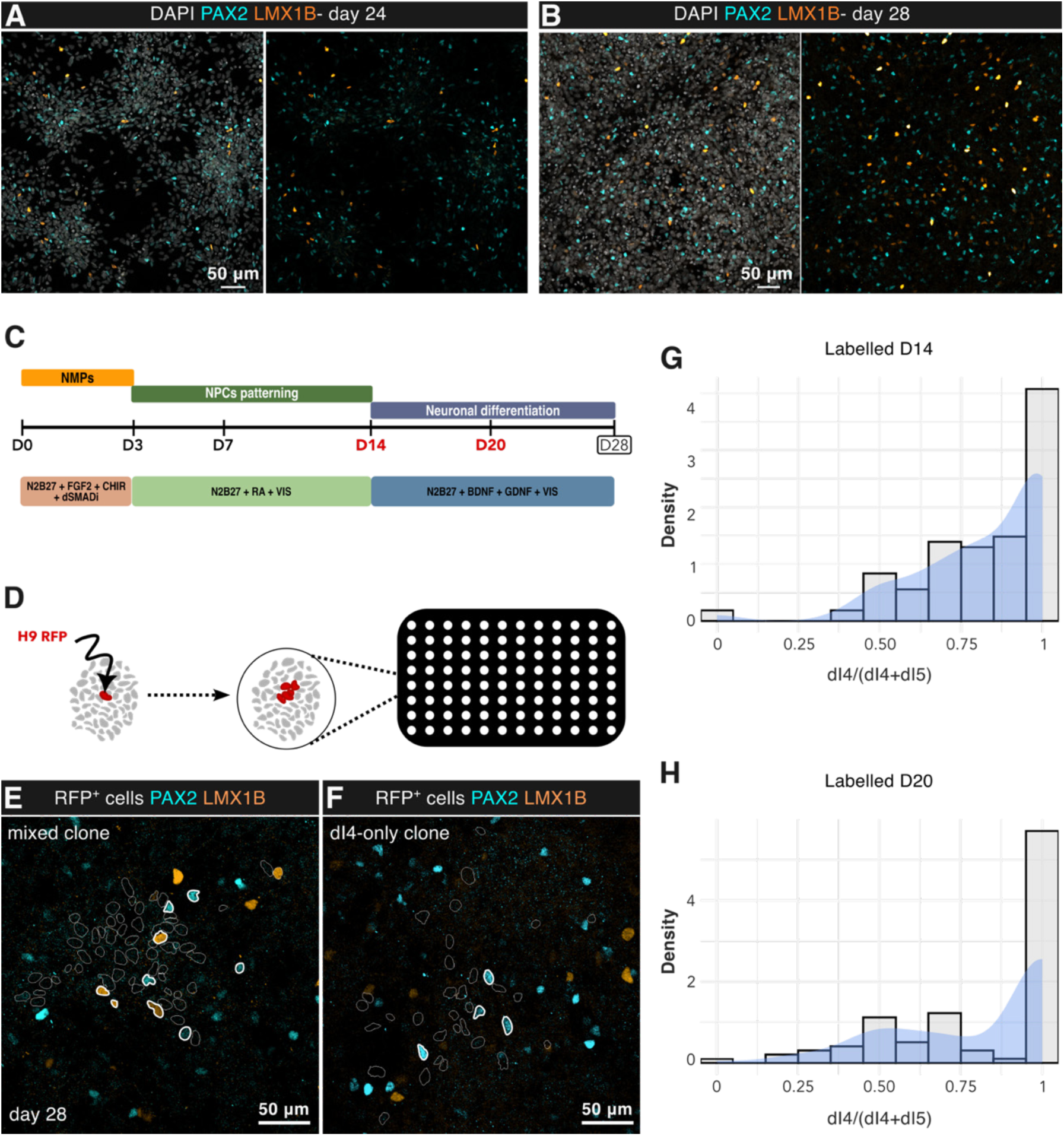
**Human stem-cell–derived model support dI4 and dI5 pedigrees** A-B. Confocal images of day 24 (A) and day 28 (B) cultures stained for PAX2 (cyan, dI4 marker) and LMX1B (orange, dI5 marker), identifying differentiating neuronal subtypes. C. Differentiation protocol of hESCs into dI4/dI5 neurons, with clonal labelling performed at day 14 and day 20 (highlighted in red) and analysed at day 28. D. Schematic of the sparse clonal labelling assay. H9-RFP cells were mixed at low density with unlabelled H9 cells in a 96-well plate to generate spatially isolated RFP-labelled clones. E-F. Confocal images of day 28 H9-RFP clones showing examples of mixed clones (E) and dI4-only clones (F). PAX2 (dI4): cyan; LMX1B (dI5): orange; H9-RFP: white. G–H. Histograms with KDE overlays of dI4 / (dI4 + dI5) ratios in clones labelled at D14 (G; 154 clones) and D20 (H; 117 clones) and analysed at D28.

To probe lineage behaviour, we sought to test the cell fate potential of clones *in vitro*. To this end, we performed clonal assays by mixing H9-RFP cells (*93*) at clonal density with unlabelled H9 cells (Fig. 5C- D, S6D-E). Progenitors were mixed at day 14, as soon as progenitor identity was established, or at day 20, capturing later progenitors closer to neuronal commitment. All were assayed at day 28 for the composition of dI4s and dI5s within clones. In both conditions, RFP⁺ clones at day 28 were either mixed (dI4 + dI5; Fig. 5E, G, H; S6F) or exclusively dI4 (dI4-only; Fig. 5F-H; S6G). dI5-only clones were virtually absent. Notably, the timing of clonal labelling affected the outcome. Early progenitors (day 14) produced mixed clones skewed towards dI4 (Fig. 5G), resembling the pattern observed in the chick HH28 dataset labelled at early stages. By contrast, later labelling (day 20) yielded mixed clones with more balanced dI4/dI5 contributions (Fig. 5H), suggesting that, while unifated dI4 progenitors persist, the later time point captures a stage when the divergence between dp4-unifated and dp4/5-bifated trajectories has already taken place.

Together, these findings are consistent with dI5s predominantly arising from bifated dI4/dI5 progenitors. Moreover, the hESC culture system provides a tractable platform for future dissection of the molecular mechanisms distinguishing dI4 sublineages and for resolving how dI4/dI5 fate decisions are balanced. Crucially, the ability to vary the timing of clonal labelling revealed stage-dependent differences in lineage output, offering a controlled experimental system to probe the time of fate commitment and how progenitor state influences fate trajectories.

### Lineage analysis of human embryos reveals conserved late bifurcation of sensory neuron fates

Having established the dual-route mechanism in chicken *in vivo* and in human ESCs *in vitro*, we asked whether it was also observed in human embryos. Although recent molecular studies have defined the timing and progression of human spinal cord neurogenesis, showing, for example, that dorsal neuron production continues beyond Carnegie stage (CS)19 (*94*), the underlying lineage relationships remain poorly understood. More recently, pseudotime analyses of human spinal cord (*95*) have proposed distinct differentiation trajectories but relied on inference and lacked direct lineage resolution. As a result, the timing and nature of lineage bifurcations between cardinal neuron classes remain unclear, especially within sensory populations, which show greater functional complexity in humans compared to other animals (*96*, *97*).

To fill this gap, we developed a new *ex vivo* slice culture system for human spinal cord explants, enabling lentiviral infection and long-term viability (Fig. 6A, S7). Transverse slices (250 μm) from CS16- CS17 embryos (6 post-conception weeks, Fig. 6A, S7A) were cultured on air–liquid interface membranes and infected with LARRY lentiviral library (Fig. 6A, S7B-C). After 7.5 days in culture, we collected barcoded cells for single-cell RNA sequencing (Fig. S6D), capturing all major spinal cord populations, including neural progenitors, neurons, and oligodendrocyte precursor cells (Fig. 6B, S6E- F). Pairwise Spearman correlation with *in vivo* reference datasets (*94*) showed the highest similarity to CS19 samples, confirming developmental progression during *ex vivo* culture (Fig. 6C). This stage corresponds approximately to HH28, the first timepoint sampled in our chick dataset (https://hdbratlas.org/comparison-HvM.html) (Fig. S8A).

**Figure 6.**
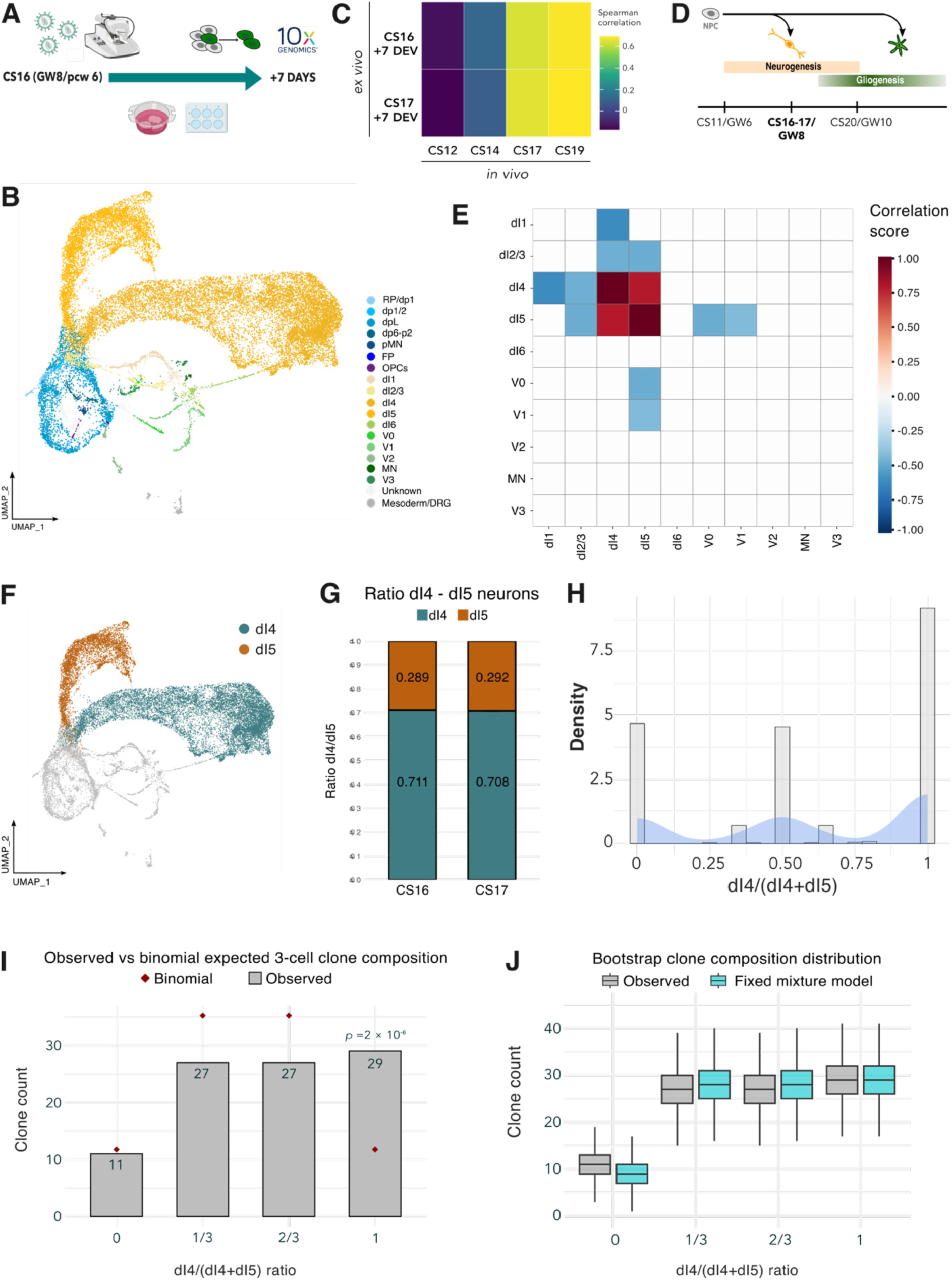
**Clonal lineage analysis in the human spinal cord** A. Diagram of LARRY barcoding experimental workflow in human *ex vivo* slices. B. UMAP of integrated human samples coloured by assigned cell types. C. Heatmap of Spearman correlation between slice cultures and *in vivo* timepoints (*92*); highest match with CS19. D. Timeline showing barcoded sample stages relative to the neurogenesis window. E. Heatmaps of neuronal clonal coupling correlations between all neuron pairs, highlighting dI4-dI5 relationships. F. UMAP visualisation of dI4 and dI5 neurons. G. Stacked bar plots showing dI4/dI5 neuron ratios in human samples. H. Histogram with overlaid kernel density estimate (KDE) of the proportion of dI4 neurons within individual clones (expressed as dI4 / [dI4 + dI5]) in human *ex vivo* samples. I. Bar plot showing the number of 3-cell clones in each dI4 / (dI4 + dI5) ratio category (0.0, 1/3, 2/3, 1.0). Red diamonds mark expected values under a binomial model assuming equal fate probability (p = 0.5, n = 3). The observed distribution significantly deviates from this expectation (exact binomial test, *p* = 2 × 10⁻⁶), primarily due to an overrepresentation of dI4-only clones. Boxplots of bootstrapped clone counts (n = 10,000 resamples) for each ratio bin under the fixed mixture model (turquoise) and observed data (grey). Strong overlap supports a good fit of the fixed model to the observed distribution.

We identified 910 multicellular clones (average multicellular clone size = 2.3 cells, max=7 cells, S8B- D). As expected, fewer multicellular clones contained cells from the ventral spinal cord compared to the chick datasets (Fig. S8E), reflecting the timing of infection in the human tissue, after most ventral neurogenesis had concluded ((*94*); Fig. 6D). As a result, labelling in the ventral neural tube predominantly targeted postmitotic neurons, yielding a high proportion of single-cell clones (93.5%).

However, lineage relationships were evident (S8E-G). For instance, dorsal neurons were linked to progenitors from their cognate domain (dI2/dI3 with dp1/3). In addition, we observed that V0, V1, dI4, dI5, dI2–3, and dI1 each contained clones with at least two neurons of the same type (Fig. S8E, diagonal line), suggesting ongoing neurogenesis within these domains. Whereas most cardinal neuronal classes showed minimal interclass clonal connections (non-diagonal values), indicating that most progenitors were committed to a domain before CS16–17, dI4s and dI5s consistently shared barcodes across both samples (Fig. 6E, S8G-H), co-occurring in 240 clones (Fig. S8E). Further supporting the equivalence of developmental stages between chick and human embryos, the dI4 to dI5 ratio (71.3% to 28.7%) matched chick HH28 proportions and was reproducible across biological replicates (Fig. 6F-G). This cross-species consistency underscores the robustness of the *ex vivo* system and suggests conserved features of dorsal interneuron patterning.

To better understand clonal dynamics, we analysed the clonal distribution of multicellular clones containing at least one dI4 or dI5. We assessed the proportion of dI4 over dI4 + dI5 cells. Similar to the chick HH28 dataset and *in vitro* experiments, we observed some clones composed exclusively of dI4 and others containing a near-equal proportion of dI4 and dI5 (Fig. 6H). To test whether the fate choice between dI4 and dI5 could be explained by independent stochastic decisions at the final cell division, as had been hypothesised for the chick (*86*), we analysed 3-cell clones, where outcomes are more interpretable. Independent fate choices (both asymmetric or randomly selected) should follow a binomial distribution, with each daughter cell independently adopting a dI4 or dI5 identity with equal probability (p = 0.5). However, the distribution of the data deviated significantly from the binomial expectation (Fig. 6I; Table 1-3), due to an overrepresentation of dI4-only clones (29 observed vs 11.75 expected; Fig. 6I). This suggests that some progenitors may have already committed to producing only dI4, rather than making stochastic choices at the last cell division. To test this, we used a mixed model that assumed the existence of both stochastic (binomial) and dI4-committed clones (Materials and Methods). This model fit the data well (χ² test, p = 0.95; Fig. 6J; Table 1, 2) and estimated that about 21% of clones came from committed dI4-only progenitors (95% CI: 11–32%; Fig. S9A), supporting the idea of a dual route to dI4s. This supports the model in which a subset of progenitors is committed to a dI4-only fate, while the remaining progenitors are bifated until the last cell division and either generate dI4s and dI5s with equal probability or in an asymmetric fashion (one dI4 and one dI5 from each division). The number of clones containing only dI5s does not exceed what would be expected under a binomial distribution (11 observed vs 11.75 expected, Table 2). In addition, the fixed mixture model fits the data well without requiring a dedicated dI5-only component, suggesting that in humans there is no detectable dp5 unifated domain, or that it is transient and already extinguished by CS16- 17, unlike the dp4-only domain that persists until later stages.

Crucially, the timing of infection and the short time between infection and collection in the human dataset provides a window onto these late fate decisions. Whereas early labelling in the chick (and in human *in vitro* at day 14) captures progenitors before bifated and unifated trajectories may diverge, making it difficult to study terminal decisions, in human embryos, the labelling likely occurs after such commitment has taken place, more similar to the *in vitro* late labelling results (day 20). This allows clearer observation of terminal fate bifurcations and strengthens the evidence for a bifated dI4/dI5 progenitor.

Overall, these findings overcome technical challenges that have limited direct lineage analysis in human tissue and indicate that, in humans, most cardinal class fate decisions are resolved by CS16, with the exception of the dI4–dI5 lineages. Occasional barcode sharing among dorsal neurons indicates that this decision window closes around six post-conception weeks. Together, these data extend our findings from the chick to human embryos, demonstrating an evolutionarily conserved feature of vertebrate spinal cord development and providing new insight into the timing and dynamics of fate specification in the human spinal cord.

More broadly, the data demonstrate how lineage tracing at different developmental stages disentangles early fate restrictions from late neurogenic decisions, offering complementary insights across species and establishing a robust framework for experimental lineage tracing in the developing human central nervous system. Together, these results not only extend our findings across species but offer a blueprint for how patterned progenitor pools shape the functional organisation of the vertebrate nervous system.

## Discussion

The assembly of functional spinal circuits depends on the orderly generation of diverse neuronal and glial cell types from molecularly distinct progenitors. While the transcriptional programs that define progenitor domains are well established, their lineage relationships have been less clear. Single-cell transcriptomics have suggested developmental trajectories (*3*, *4*, *95*), yet without direct clonal validation, these models remain provisional and are often confounded by processes such as phenotypic convergence or divergence. Here, by combining genomic barcoding with single-cell transcriptomics in chick and human, we reconstruct progenitor pedigrees at cell-type resolution. Our analysis reveals a previously unappreciated hierarchical organisation of progenitor domains that aligns with circuit-level function and identifies developmental trajectories for neuronal subtype specification. Together, these findings suggest that lineage compartmentalisation is an organising principle that links the evolutionary origin of cell type diversification to embryonic patterning and circuit assembly.

### Lineage subdivisions link spatial patterning to functional circuit assembly

Early retroviral tracing in the chick suggested broad and relatively uniform progenitor dispersal along the dorsoventral axis (*35–37*), consistent with a continuum of lineage relationships. By contrast, molecular models of spinal cord patterning have defined eleven progenitor domains as independent, fate-restricted units defined by their transcription factor codes (*11*, *98*, *99*). Our clonal analyses refine these perspectives. Rather than uniform mixing or strictly isolated domains, progenitors clustered into five modular lineage subdivisions that sit hierarchically above the 11 canonical progenitor domains. The domains themselves were evident, with progenitors most often producing neurons within their own domain, but lineage relationships extended across neighbouring domains. These subdivisions capture shared developmental origins across neighbouring domains while constraining lineage relationships within defined groups. Support for these discrete subdivisions also comes from earlier genetic labelling studies in mouse, which showed descendants of Dbx1-Cre⁺ remaining largely confined within a restricted dorsoventral region, resembling the spatial boundaries of our proposed lineage subdivisions (*100*).

The hierarchical organisation of lineage subdivisions suggests a mechanism establishing progenitor domain patterns along the dorsal-ventral axis. Rather than emerging through linear, sequential induction by graded morphogens acting at a series of monotonic thresholds, our findings indicate that the patterning cues control bifurcating binary decisions that progressively refine progenitor commitment. In this model, cells first select between broader lineage subdivisions, then resolve into specific progenitor domains within each subdivision (Fig. 2D). This is consistent with recent dynamical systems modelling of spinal cord development, which suggests that progenitor diversification is achieved through a series of discrete binary bifurcations (*54*). The most prominent subdivision boundary between dI5 and dI6 exemplifies this logic: despite being spatially adjacent, these populations arise from distinct lineage branches, suggesting that this boundary reflects a fundamental bifurcation point rather than a simple morphogen threshold. This modular structure aligns with molecular evidence: chromatin accessibility profiles distinguish p3/floor plate progenitors from those in pMN and p2, which share regulatory landscapes (*53*), reinforcing the notion that lineage architecture mirrors underlying gene regulatory logic.

The lineage subdivisions are aligned with neuronal function. For example, dI6 interneurons, conventionally classified as dorsal (*55*, *59*), instead clustered with ventral V0–V1 interneurons, consistent with their shared function in locomotor circuits (*101*). This is in line with some pseudotime analyses (*6*), while it diverges from others (*95*), underscoring the necessity of direct lineage reconstruction. Our clonal data provide this direct *in vivo* evidence, revealing that lineage subdivisions not only reflect spatial patterning but also anticipate functional circuit organisation, linking developmental logic with emergent neural function.

### Temporal competence shifts generate diversity within spatial lineage compartments

Clonal analysis revealed that single progenitors generate neurons from different temporal waves, indicating that individual progenitors undergo competence shifts rather than being restricted to a single neurogenic window. This distinction is important: while temporal waves of neurogenesis in vertebrates are well established (*2*, *26–28*, *102*), it has remained unclear whether each wave arises from distinct progenitor pools or whether single progenitors change their output over time. Our data support the latter, paralleling Drosophila neuroblasts (*103–108*). Similarly, in the cortex, individual radial glia progenitors undergo progressive competence shift, giving rise to neurons in several layers (*109–111*). While temporal output shifted over time, spatial identity remained stable across subdivisions. Individual progenitors generated neurons of different cardinal classes across successive waves. In the ventral spinal cord, we observed a subtle ventral bias, with mid-wave neurons often clonally linked to later-born neurons from more ventral domains. This is consistent with the idea of kinematic waves of gene expression progressively ventralising progenitor identity over time (*8*). As an example, Nkx2.2⁺ p3 progenitors, associated with the generation of V3 interneurons, have been shown to transiently generate motor neurons (*31*). Supporting this, our clonal data reveal late p3 links to MNs, consistent with progressive ventralisation of the pMN lineage. We also find that oligodendrocyte progenitors (OPCs) are related to neurons from different domains, reconciling conflicting reports of pMN-exclusive (*33*, *112–114*) versus p3-contributing origins (*31*, *81–83*). It will be important to assess whether a similar progressive model applies to dorsal progenitors influenced by roof plate signals. Together, these observations support a model in which spatially defined progenitors undergo sequential competence shifts to generate diverse neuronal and glial lineages.

### Dual developmental routes and functional specialisation of sensory interneurons

In the dorsal spinal cord, pseudotime trajectories, gene expression and knock-out studies have proposed that dorsal interneurons arise either through two lineage groups (dI1-3 and dI4-6) or alternating sets (dI1-3-5 and dI2-4-6) (*6*, *21*, *22*, *55*, *60*). Our clonal data rule out the alternating model and instead support two dorsal lineage groups: dI1–2 and dI3–5. The data also support the idea that dI4 and dI5 sensory interneurons are generated through distinct developmental routes. Fast differentiating unifated dp4 progenitors give rise exclusively to dI4s, whereas slower differentiating bifated dp4/5 progenitors produce both dI4 and dI5. This organisation is consistent with the rarity of dp5-only progenitors and initial lower number of dI5s. While we cannot exclude a transient, dedicated dp5 lineage, as suggested by previous studies (*55*, *58*, *85*, *86*), its contribution would be minimal, either exhausting rapidly or transitioning into bifated output. These findings refine the canonical dp4/dp5 model (*55*, *58*, *85*, *86*), suggesting that unifated dp4 progenitors may contribute more substantially and persist longer than dp5 progenitors, with bifated dp4/5 progenitors contributing to the majority of sensory neurons due to their delayed output. Further analyses across developmental time points, incorporating spatial information, will be needed to clarify the contribution of dp5 progenitors. In this context, it will be important to assess the dynamics that give rise to dI3 neurons, which appear to have more flexible links with dp1–2 progenitors as well as with more ventral dp4/5 and dI4 neurons, possibly reflecting distinct temporal patterns of progenitor specification.

Notably, the origin of dI4s and dI5s parallels other systems where progenitor plasticity is retained until the final division, generating both excitatory and inhibitory neurons (*115–117*). Notch signalling is a common mediator of such terminal decisions (*38*, *51*, *116*, *118*). Although direct evidence remains limited, a role for Notch in the dI4/dI5 fate decision has been proposed (*85*). The consequences of such a terminal bifurcation may be functionally relevant to balance excitatory and inhibitory output by buffering variability derived from progenitor proliferation or differentiation noise. Indeed, loss or reduction of *PTF1A* expression in mice and humans induces severe circuit imbalance manifesting with increased itch sensitivity (*57*, *119*) or irregular respiration and cerebral hyperexcitability (*120*). Together, these findings underscore the physiological importance of balancing inhibitory and excitatory neurons, suggesting that terminal divisions may act as a safeguard for circuit homeostasis in pain, itch, and touch circuits.

Beyond balancing excitation and inhibition, lineage history also appears to shape molecular and functional specialisation within the dI4 class itself. dI4 arising from unifated clones expressed higher *TFAP2B*, consistent with motor-synergy encoder identity (*87*) and deep dorsal horn localisation (*5*), whereas those from bifated clones expressed elevated *NPY, PRKCA/D* and *KCNIP4*, marking mechanical-itch gating neurons located in the superficial dorsal horn (*88*, *89*).

### Evolutionary origins and comparative CNS organisation

While transcriptomic and pseudotime analyses have suggested lineage relationships in the human spinal cord (*94*, *95*), direct clonal evidence has been lacking due to the difficulty of *in vivo* manipulation. Lineage tracing at this resolution in human tissue opens a window into previously inaccessible stages of development. Our data show that most fate commitment between progenitor domains is already resolved by around six weeks post-conception, marking a key developmental milestone. Within the dorsal spinal cord, where sensory populations continue to expand, we find that the dual-route strategy identified in chick is conserved in human *ex vivo* and *in vitro,* suggesting a conserved basis for sensory interneuron diversification. The similarity between chick and human underscores the value of the chick spinal cord as a model to study lineage organisation in vertebrate neurogenesis.

Lineage studies suggest that the brain, like the spinal cord, is organised into discrete clonal compartments. Retroviral and genetic lineage tracing first distinguished pallial and subpallial origins in the telencephalon (*121–129*) and high-throughput genomic barcoding has confirmed and refined this view (*45*). Within these broad compartments, a nested hierarchical organisation can be appreciated. The medial (MGE) and lateral (LGE) ganglionic eminences, both subpallial derivatives, generate largely separate neuronal lineages (*125*, *127*) and exhibit internal domain-specific biases (*130–132*). Thus, as in the spinal cord, domains in the forebrain appear nested within broader clonal subdivisions, arguing that regional lineage compartmentalisation is a general organisational principle of the central nervous system (CNS). This conserved logic also raises the question of its evolutionary origin.

The compartmentalisation of the spinal cord highlighted by our data is reminiscent of the ancestral bilaterian “columnar” plan, in which dorsal, intermediate, and ventral stripes are defined by the expression of the *Drosophila* transcription factors msh/Msx, ind/Gsh, and vnd/Nkx2, respectively (reviewed in (*133*)). Consistent with this framework, amphioxus also expresses Msx and Nkx2/6 orthologues in ordered dorsoventral domains (*134*, *135*), and annelids show similar DV patterning (*136*, *137*). These parallels suggest that vertebrate progenitor architecture may have evolved from an ancient tripartite scaffold, subsequently diversified by further subdivisions (*138*).

In non-vertebrate bilaterians, a ventral compartment equivalent to dI6–V1 appears to be absent. Neurons from this lineage innervate motor neurons, coordinating and modulating gait (*64*, *65*, *101*, *139*). The emergence of this compartment may therefore represent a vertebrate elaboration on the ancestral scaffold, supporting increasingly sophisticated motor control systems (*101*, *140*). Consistent with this, lamprey embryos express *Dbx1* in the central spinal cord (*141*), flanked by ventral *Nkx2-2* and dorsal *Pax3/7* (*142*, *143*), hinting that this compartment may have emerged in jawless vertebrates. Notably, the telencephalon lacks an analogous intermediate progenitor domain: within the subpallium, gene expression distinguishes NKX2-2⁺ MGE from GSX2-enriched LGE (*144*, *145*), but no discrete intermediate domain separates them. Thus, while the spinal cord evolved a dedicated dI6– V1 lineage module, the telencephalon retained a simpler bipartite organisation. This suggests that the addition of this intermediate inhibitory subdivision was a specific innovation of the vertebrate spinal cord, tied to the evolutionary demands of locomotor coordination (*146*). In this light, the nested lineage compartments could reflect a conserved bilaterian architecture expanded by vertebrate-specific additions. The incorporation of such modules may have enabled the diversification and patterning of motor circuits that define the vertebrate clade.

## Limitations and future directions

While the approach establishes a framework for molecularly annotated reconstruction of lineages, barcode dropout means that only a sample of cells in a clone are recovered. This sets an upper limit on the power of modelling predictions and prevents definitive conclusions when no clonal sharing is observed. Moreover, while clonal outputs could be linked to mature cell identity, the current barcoding methods lack spatial data and molecular information about progenitors at the time of labelling, making it impossible to conclusively connect competence shifts or lineage restrictions with position or progenitor gene expression. Addressing these limitations will require methodological advances that integrate lineage tracing with spatial readouts and concurrent detailed molecular profiling.

Looking forward, our work raises several biological questions. What molecular mechanisms, transcriptional programmes, morphogen gradients, and adhesion codes implement the lineage subdivisions? How do temporal waves in progenitors couple to the birth order of neuronal cohorts? Is the dI4/dI5 fate decision stochastic, or deterministic and asymmetric? How is the dI4/dI5 bifurcation implemented? What are the temporal progressive lineage restrictions within subdivisions? Finally, to what extent are the modular principles revealed here generalised across the CNS? Answering these questions will require combining clonal recording with simultaneous progenitor profiling, perturbations and tractable *in vitro* systems. The framework we provide offers a foundation for this next phase and a path to connect embryonic patterning with circuit assembly.

## Materials and Methods

### Library amplification

The LARRY Barcode Version 1 plasmid library (Addgene #140024), a gift from Fernando Camargo, was amplified for downstream applications. A total of 20 ng of plasmid DNA was electroporated into each of 12 vials of Stbl4 ElectroMAX™ cells (25 µL per vial; Thermo Fisher Scientific, Cat. No. 11635018). Following a 1-hour recovery at 37°C in SOC medium, each vial was plated onto two 20 × 20 cm LB-agar plates. The following day, bacterial colonies were rinsed from the plates using a total of 1.5 L of LB medium, which was then cultured for 2.5 hours at 37°C. Plasmid DNA was purified using the NucleoBond® Xtra Maxi EF kit (Macherey-Nagel, Cat. No. 740424.50), yielding 1–1.5 mg of DNA.

### Lentivirus production

LARRY lentivirus was generated using a third-generation packaging system comprising pLP1 and pLP2 (Invitrogen, Cat. No. A43237), and pVSV-G (System Biosciences). HEK293T cells were cultured in DMEM (Sigma, Cat. No. D5796) supplemented with 10% fetal bovine serum (FBS, Biosera, Cat. No. FB- 1001/500) and 1% penicillin-streptomycin (Gibco, Cat. No. 15140-122). Transfection was carried out in Opti-MEM™ I (Gibco, Cat. No. 31985) using Lipofectamine 2000 (Invitrogen, Cat. No. 11668500). Sixteen hours post-transfection, the medium was replaced with fresh DMEM supplemented as above. Viral supernatants were collected at 36- and 60-hours post-transfection, filtered through a 0.45 μm filter (Millex-HV, Millipore, Cat. No. SLHV033RS) and ultracentrifuged at 22,000 rpm for 1.5 hours. Viral pellets were resuspended in sterile DPBS (Gibco, Cat. No. 14190) and stored at −80 °C. Viral titres ranged from 1 × 10⁸ to 1 × 10⁹ TU/mL.

### Plasmid and lentiviral library diversity assessment

To assess library diversity after expansion, we sequenced the barcoded region of both plasmid and lentiviral libraries. For the plasmid library, barcodes were amplified using primers: pLARRYfw: TCGTCGGCAGCGTCAGATGTGTATAAGAGACAGaggaaaggacagtgggagtg and pLARRYrv: GTCTCGTGGGCTCGGAGATGTGTATAAGAGACAGcaaagaccccaacgagaagc. For the lentiviral library, RNA was extracted from 40 µL of virus using the NucleoSpin® RNA Virus kit (Macherey-Nagel, Cat. No. 740983.50), and eluted in 40 µL. 3 µg of RNA were reverse transcribed to cDNA using SuperScript™ VILO™ (Thermo Fisher, Cat. No. 11754050) with random hexamers. Amplification was performed using vLARRYfw: TCGTCGGCAGCGTCAGATGTGTATAAGAGACAGaatcctcccccttgctgtcc and vLARRYrv: GTCTCGTGGGCTCGGAGATGTGTATAAGAGACAGaccgttgctaggagagaccata. To prevent saturation, cycle numbers were determined via mock qPCR. Amplification was performed with Q5® High-Fidelity DNA Polymerase (NEB, Cat. No. M0494) under the following conditions: 98°C for 30 s; [98°C for 10 s, 63°C for 30 s, 72°C for 20 s] for 7–15 cycles; final extension at 72°C for 2 min. Libraries were dual-size selected (0.5X–0.2X) using AMPure XP beads (Beckman Coulter, Cat. No. A63881) and sequenced to ∼5 million reads (plasmid) or ∼10 million reads (virus) on a MiSeq and NovaSeq 6000 platform, respectively.

Barcode diversity was assessed using the genBaRcode R package (v1.2.5) (*147*), identifying > 250,000 unique barcodes in both libraries, consistent with the originally reported diversity (*43*).

### Chicken embryos and injections

Fertilised chicken eggs (*Gallus gallus*) were obtained from Henry Stewart & Co. Ltd. and incubated at 38°C in a humidified incubator until the desired Hamburger–Hamilton (HH) stage. HH11–HH12 embryos (40–48 hours incubation) were injected into the neural tube canal from the anterior end with concentrated lentivirus (1 × 10⁸ – 1 × 10⁹ TU/mL) mixed with SYBR™ Green (Thermo Fisher, Cat. No. S7563) using pulled glass capillaries (Harvard Apparatus, Cat. No. EC1 64-0766). Embryos were harvested at stages HH28 (day 6), HH31 (day 7.5), or HH35 (day 9). All embryos were cultured for no more than two-thirds of their development.

### Human embryos

Human embryonic material was provided by the MRC/Wellcome Human Developmental Biology Resource (grant MR/R006237/1; https://www.hdbr.org), with appropriate maternal written consent and approval from the London Fulham Research Ethics Committee (18/LO/0822) and the Newcastle and North Tyneside NHS Health Authority Joint Ethics Committee (18/NE/0290). HDBR is regulated by the UK Human Tissue Authority (HTA; www.hta.gov.uk) and operates in accordance with the relevant HTA Codes of Practice. This work was part of project no. 200668 registered with the HDBR.

The HDBR provided fresh tissue from embryos and fetuses aged 8–12 gestational weeks (GW; 6-10 post conception weeks -pcw-), corresponding to the following Carnegie stages (CS): CS16 (n = 2), CS17/18 (n = 2), CS21 (n = 1), CS23 (n = 2), 9 pcw (n = 1), and 10 pcw (n = 2). In accordance with data protection policies, information on the sex or gender identity of the embryos/fetuses is not available. These factors are not expected to influence the results reported in this study. Spinal cord tissue was dissected in L-15 medium and either used immediately for slice culture or fixed for downstream applications. Samples that were not suitable for lentiviral infection—due to insufficient material or an advanced developmental stage—were used for slice culture optimisation or fixed for immunohistochemistry. Fixation was performed in 4% paraformaldehyde (PFA) in 0.12 M phosphate buffer (pH 7.4) overnight at 4°C.

### Human organotypic slice cultures

Trunk tissue was dissected in cold L-15 medium by removing surrounding non-neural tissue, then embedded in 4% low-melting-point agarose (Sigma, Cat. No. A2790). Transverse spinal cord slices (250 μm thick) were prepared using a vibratome (Leica VT 1200S). Intact slices were selected and transferred to 6-well plates containing cell culture inserts (0.4 μm pore) for air–liquid interface culture (Sarstedt, Cat. No. 83.3930.040). Each well contained 1.5 mL of slice culture medium (SCM). SCM was prepared as described by Long et al. (*148*) and consisted of Neurobasal medium (Gibco, Cat. No. 21103049) supplemented with 10% KnockOut™ Serum Replacement (KOSR; Gibco, cat. 10828010), 1X GlutaMAX™ (100X stock, Gibco, cat 35050061), 1X Penicillin–Streptomycin (100X stock, Gibco, cat. 15140122), 1X N-2 Supplement (100X stock; Gibco, 17502048), 1X B-27 Supplement (50X stock; Gibco, 17504044), and 0.1 M HEPES-NaOH (Sigma, Cat. SRE0065), pH 7.2. A drop of 2-5 μL of concentrated LARRY lentivirus (1 × 10⁸ to 1 × 10⁹ TU/mL), mixed with SYBR™ Green (Thermo Fisher, Cat. No. S7563), was applied onto the slice into the central canal of the neural tube using a P10 pipette tip. Slices were maintained in a tissue culture incubator under 20% O₂ / 5% CO₂ at 37°C for 7.5 days. Half-medium changes were performed every other day. Slices were inspected regularly for signs of necrosis, and only healthy-appearing slices were selected for downstream single-cell RNA sequencing.

### Single-cell dissociation and flow cytometry

For chick samples, brachial/thoracic spinal cords were freshly dissected at stages HH28 (n = 4), HH31 (n = 7), and HH35 (n = 6) in cold L-15 medium (Gibco, Cat. No. 15460554). For human samples, single-cell dissociation was performed from previously cultured spinal cord slice preparations (see above). Each human sample consisted of 10–12 healthy slices manually detached from membrane inserts using an eyelash manipulator (Clinisciences Limited, cat. 71182). Two embryos (CS16 and CS17) were processed.

Tissues were enzymatically dissociated in FACSmax dissociation buffer (amsbio, cat. AMS.T200100) supplemented with 1mg/ml papain (Sigma, cat.10108014001) at 37°C for 20 minutes with shaking at 700 rpm, and occasional gentle pipetting. Cells were pelleted at 0.6 rcf for 4 minutes, resuspended in HBSS supplemented with 1% BSA, and filtered through a Flowmi® 40 µm cell strainer (Sigma, cat. BAH136800040-50EA) to obtain a single-cell suspension. Cells were transferred to round-bottom FACS tubes and stained with 1 µg/mL DAPI for viability assessment.

Fluorescence-activated cell sorting was performed with a BD FACSAria™ Fusion or BD Influx™ with a 100 μm nozzle at 4°C. Viable GFP⁺/DAPI⁻ cells, as well as GFP⁻/DAPI⁻ control cells, were sorted into DNA LoBind tubes (Eppendorf, Cat. No. 1070870) pre-coated with HBSS containing 1% BSA. Cell concentration and viability were assessed with a LUNA™ automated counter (Logos Biosystems). Cells were pelleted at 0.5 rcf for 5 minutes at 4°C, the supernatant was removed, and cells were resuspended in PBS with 0.04% BSA to the appropriate concentration for downstream applications.

### Single-cell RNA sequencing and cell type annotation

Single-cell libraries were prepared using the Chromium Next GEM Single Cell 3ʹ Reagent Kit v3.1 (10x Genomics; for chicken samples) or v4 (10X Genomics; for human samples) following the manufacturer’s protocol. Quality control of cDNA and final libraries was performed using the 4200 TapeStation System (Agilent Technologies) and Qubit dsDNA HS Assay Kit (Thermo Fisher). Sequencing was carried out on NovaSeq 6000/NovaSeq X platforms using a 28–10–10–90 read setup.

Raw FASTQ files were processed with Cell Ranger 9.0.0 against the *Gallus gallus* GRCg7b or human GRCh38 genome assemblies. An additional contig comprising of the GFP and LARRY sequence were added to the assemblies to quantify the GFP and LARRY expression. Initial quality control in Scanpy retained cells with sufficient transcriptome complexity (>2,000 detected genes) and excluded those with abnormally high library size (>45,000 UMIs). Putative doublets were identified with Scrublet and removed (*149*). For initial integration and clustering, we selected 2,000 highly variable genes (HVGs) with Scanpy and trained a variational autoencoder with scVI (n_latent = 30). After the first scVI run, clusters corresponding to non-neural contaminants (e.g., blood, mesoderm, neural crest/DRG) were removed before repeating the HVG selection and scVI training on the cleaned dataset. The HVGs capture genes with high variability across cells, which is useful for batch correction and latent embedding, but they can also include noise-driven genes. To refine the feature space, we therefore applied continuous Entropy Sort Feature Weighting (cESFW;(*150*, *151*). This method identifies genes that are structured and co-expressed across the dataset, highlighting biologically informative features while filtering out uninformative noise. After visual inspection of the cESFW gene–UMAP, we retained the around 2,000 weighted genes (the choice was performed separately for each dataset) and retrained scVI on this subset.

The final cESFW-informed scVI latent representation was used for nearest-neighbours graph construction, UMAP embedding, and Leiden clustering across a range of resolutions. Data were intentionally overclustered to ensure separation of transcriptionally distinct populations. Cluster identities were then annotated based on established neural progenitor and neuronal subtype markers (Table 4, Fig. S1G).

### LARRY library amplification

To enrich for LARRY barcodes, we separately amplified the barcoded region from cDNA generated during the Chromium single-cell library preparation and sequenced it independently from the gene expression (GEX) library. Amplification was performed using Q5® High-Fidelity DNA Polymerase (NEB, Cat. No. M0494) in 50 µL reactions containing 1X Q5 2× Master Mix, 0.3 µM i7-indexed reverse primer, 0.3 µM i5-indexed forward primer, molecular-grade H₂O, and 4 µL of cDNA template (for primer sequences and indices, see Table 5). PCR cycling conditions were as follows: initial denaturation at 98 °C for 30 s; followed by 14–19 cycles of 98 °C for 10 s, 64 °C for 30 s, and 72 °C for 30 s; and a final extension at 72 °C for 2 min. Libraries were purified using dual-sided size selection (0.5X–0.2X) with AMPure XP beads (Beckman Coulter, Cat. No. A63881) and quantified using an Agilent BioAnalyzer or TapeStation. Enriched LARRY libraries were sequenced on NovaSeq 6000/NovaSeq X platforms using a 28–10–10–90 read setup, to a depth of approximately 5,000 reads per cell, representing ∼10% of the transcriptome library sequencing depth.

### Clonal assignment and barcode filtering

Cells carrying LARRY barcodes were assigned to clones using a custom Nextflow pipeline pipeline (available at https://github.com/FrancisCrickInstitute/LARRY), inspired by Weinreb et al (*43*). For each sample, both scRNA-seq data and the corresponding amplified LARRY libraries were processed. Valid cell barcodes were identified using the cellranger count module from nf-core (*152*). Adapter sequences were removed from the LARRY libraries using the cutadapt module (*152*, *153*), and only reads with valid cell barcodes, UMIs, and LARRY barcodes were retained. UMIs were extracted and appended to read names using the umitools extract module(*152*, *154*), using the barcodes from cellranger as a whitelist. A custom script was then used to extract cell barcode, UMI, and LARRY barcode information, and to quantify LARRY barcode expression per cell, based on UMI counts. To distinguish background from true barcode expression (e.g., ambient RNA), a per-sample expression threshold was determined: LARRY barcodes were iteratively removed starting from the lowest UMI counts, decrementing counts by one at each step. The process stopped when the mean UMI count stabilised (relative change ≤ 0.01), and this point was used as the expression cutoff. To account for multiple infections, cells could be assigned multiple LARRY barcodes. Clones with Jaccard similarity ≥ 0.5 were merged. Cells that remained associated with multiple clones after merging were excluded from downstream clonal analyses.

### Permutation testing and clonal co-occurrence analysis

To assess which cell types tend to co-occur within clones, we calculated clonal coupling z-scores (*44*, *46*, *50*). To establish a baseline for expected co-occurrence, we performed 1,000 random permutations in which the number of clones, clone sizes, and the number of cells of each type were preserved. In each permutation, cells were randomly assigned to clones, and co-occurrence frequencies were computed for all cell type pairs. We compared the observed co-occurrence frequencies to this null distribution. For each cell type pair, we calculated a z-score reflecting how many standard deviations the observed frequency was from the permuted mean, with positive z-scores indicating more frequent co-occurrence than expected by chance, and negative scores indicating the opposite. To explore broader patterns of lineage relationships, we computed a Pearson correlation matrix of these z-score profiles (*46*, *50*). For each cell type pair, this correlation measures how similarly they co-occur with all other cell types. For visualisation, we only plotted correlation values where the original z-score and the correlation coefficient had the same sign: either both positive (shared lineage) or both negative (mutual exclusivity). When the signs differed, the matrix cell was left blank to avoid displaying misleading associations. To explore higher-order structure in the correlation matrix, we performed hierarchical clustering using Ward’s method on the pairwise Pearson distances (1 - correlation coefficient). This approach groups cell types that share similar clonal coupling profiles, revealing clusters with common patterns of co-occurrence or mutual exclusivity. The resulting dendrogram provides a lineage-informed organisation of cell types, in which proximity reflects shared clonal behaviour across the dataset.

### Correlation of *in vivo* and *ex vivo* single-cell data

To compare *ex vivo* slice cultures with in vivo spinal cord developmental stages, we identified a subset of 781 genes that best separated cell states in the *in vivo* dataset (*94*) using entropy sorting gene selection. For each *in vivo* time point, pseudo-bulk gene expression profiles were generated by averaging the expression values of these genes across all cells. The same procedure was applied to each *ex vivo* slice sample. Spearman correlation coefficients were then calculated between the resulting pseudo-bulk profiles to assess similarity between *ex vivo* samples and *in vivo* time points.

### Modelling clone composition of human dI4/dI5 sensory neurons

To evaluate whether the distribution of dI4/dI5s in 3-cell clones could be explained by stochastic fate decisions, we compared the observed clone composition to a binomial model assuming equal probabilities (p = 0.5) at the final division. Expected clone counts were computed for each composition bin (0/3, 1/3, 2/3, 3/3 dI4), and goodness-of-fit was assessed using χ² tests and exact binomial tests.

To account for potential lineage bias, we fitted two mixture models. The fixed mixture model combined a binomial component (parameterised by α) and a dI4-only committed component (1 – α), with α estimated by minimising the χ² distance between observed and expected distributions. Confidence intervals for α were computed using profile likelihood.

Model fit was evaluated using the log-likelihood, Akaike Information Criterion (AIC), and Bayesian Information Criterion (BIC), both of which penalise overfitting by accounting for model complexity (Table 3). The fixed mixture model showed substantially better AIC and BIC scores compared to the binomial model, indicating a more parsimonious and accurate fit despite the added complexity. To further assess model robustness, we generated 10,000 bootstrap replicates of the observed and model-predicted clone distributions. For both the model and the observed data, we simulated new datasets by sampling clone ratios according to their probability distribution, preserving the original clone count. For each replicate, we computed the χ² statistic, yielding empirical null distributions used to derive bootstrap-based p-values as the proportion of χ² values greater than or equal to the observed χ² (Table 1). We also computed the mean and 95% confidence intervals for each bin across bootstraps and compared them to the observed values (Table 2), visualised as boxplots (Fig. 6J). Additionally, histograms of the bootstrapped χ² distributions, with vertical lines marking the observed χ² values, facilitated direct comparison across models (Fig. S9B). All statistical analyses were performed in R, and significance was evaluated at a threshold of p < 0.05.

### Immunohistochemistry of embryo sections

Chicken and human embryos and human slice cultures were fixed in 4% paraformaldehyde for 2 h at room temperature (chicken) or overnight at 4 °C (human), then cryoprotected overnight at 4°C in 0.12 M sodium phosphate buffer with 15% (w/v) sucrose. Samples were embedded in phosphate buffer with 15% sucrose and 7.5% gelatin, then flash-frozen in isopentane. Cryosectioning (14 μm) was performed using a Leica CM3050S cryostat onto SuperFrost slides (Thermo Fisher, Cat. No. 10149870). Sections were stained following standard protocols (*90*). Primary antibodies included: mouse anti- NKX6.1 (1:100, DSHB F55A10), goat anti-OLIG2 (1:500, R&D Systems AF2418), rat anti-SOX2 (1:500, Santa Cruz sc-365823), rat anti-RFP (1:500, Chromotek 5F8), rabbit-Tubb3, goat-SP8, mouse-NKX2-2, mouse PAX7, rabbit HuC/D, rabbit PAX6, sheep-GFP, rabbit-PAX2, guinea pig-LMX1B, guinea-pig LBX1, mouse-MSX1, mouse-OLIG3, rabbit-SOX10, mouse LHX1, goat ISL1 (1:500, R&D, AF1837), Casp3 (??). Alexa Fluor-conjugated secondary antibodies (Thermo Fisher) were used at 1:500 dilution. DAPI (Sigma, Cat. No. MBD0015, 10ug/ml) was used for nuclear counterstaining. Imaging was performed using a Leica SP8 confocal microscope, and images were processed with Fiji or Imaris.

### Whole-mount immunostaining of human slice cultures

At the end of the culture period, slice cultures were embedded in low-melting-point agarose on glass-bottom dishes and fixed in 4% paraformaldehyde (PFA) in PBS for 1 h at 4°C, followed by three washes in PBS. Permeabilisation and blocking were carried out for 1–2 h at room temperature in PBS containing 0.2% gelatin and 0.5% Triton X-100 (PBSGT). Samples were then incubated with primary antibodies diluted in PBSGT supplemented with 0.1% saponin (1 mg/mL) for 48 h at 4°C. After six washes in PBSGT at room temperature, slices were incubated for an additional 48 h at 4°C with secondary antibodies and DAPI (10 µg/mL) diluted in PBSGT containing 0.1% saponin (1 mg/mL). Samples were washed six times in PBSGT (30 min each), followed by three final washes in PBS, and stored in PBS at 4°C for short-term preservation prior to imaging.

Primary antibodies used were rat anti-SOX2 (1:500, Santa Cruz sc-365823), rabbit-Tubb3sheep-GFP. Alexa Fluor-conjugated secondary antibodies (Thermo Fisher) were used at 1:500 dilution.

### Clonal size estimation from chick neural tube sections

Clonal size was assessed from transverse sections of embryos injected at HH12 and collected at HH28, a developmental stage at which clones remained relatively compact and could be reliably identified. Only sections containing a maximum of two well-separated clones were included in the analysis to avoid overlap and misidentification. LARRY⁺/DAPI⁺ cells were manually counted for each clone. Dorsal and ventral clones were classified based on their position relative to the forming sulcus limitans; clones with ambiguous localisation were excluded from the analysis.

### Directed differentiation of hESCs into dI4/dI5 fates

H9 ESC line (WiCell), H9-RFP ESC (*93*) lines were routinely cultured in Stemflex medium (Gibco, A3349401) on 0.5-μg/cm2 laminin-coated plates (Thermo Fisher Scientific A29249) and split using ReLeSR™ (Catalog # 100-0483, StemCell Technologies).

Human embryonic stem cells (hESCs) were directed toward dorsal interneuron (dI4/dI5) lineages using a protocol adapted from previously published methods (*60*, *90–92*) (Fig. 5E). Cells were seeded on vitronectin-coated plates (Cat. No. A14700; 1:100 in DPBS) at a density of 60,000 cells/cm² and cultured in N2B27 medium supplemented with 20 ng/mL FGF2 (Cat. No. 100-18B), 3 µM CHIR99021 (Axon, Cat. No. 1386), 2 µM DMH1 (Cat. No. A12820), 10 µM SB431542 (Cat. No. HY-10431), and 10 µM Y-27632 (Tocris, Cat. No. 1254)was included for 24 hours. From day 1 to day 3, the same medium was used without Y-27632. On day 3, cells were replated onto Geltrex-coated plates (Cat. No. A1413302) and cultured in N2B27 supplemented with retinoic acid (RA, Cat. No. R2526; 100 nM), Vismodegib (GDC-0449; APExBIO; 0.5 µM), and Y-27632 for 24 hours. From day 4 onward, cells were maintained in RA and Vismodegib without Y-27632. A second replating onto Geltrex-coated coverslips was performed on day 7 at 80,000 cells/cm². From day 8 to day 14, cells were cultured in N2B27 with RA and Vismodegib. A final replating was performed on day 14 at 100,000–150,000 cells/cm² in a medium containing Vismodegib (0.5 µM), brain-derived neurotrophic factor (BDNF, Cat. No. 17874523), glial cell line-derived neurotrophic factor (GDNF, Cat. No. 17874303), both at 20 µg/mL, and Y-27632. From day 15 onward, media were refreshed every 2–3 days with N2B27 containing Vismodegib, BDNF, and GDNF. Vismodegib was reduced to 0.2 µM from day 22, and cells were fixed between days 24 and 28 for downstream analyses.

### Clonal analysis of differentiated hESC-derived interneurons

For clonal analyses, H9 and H9-RFP cells were co-cultured by mixing a high-density population of H9 cells (125000 cells/cm2) with a sparse number of H9-RFP cells (Fig. 5F). On day 14 and day 20 of differentiation, H9 cells were dissociated and replated onto Geltrex-coated 96-well plates (Revvity PhenoPlate 96, Cat. No. 6055302) at 125,000 cells/cm² (day 14) or 150,000 cells/cm² (day 20), using the same medium conditions described above. H9-RFP cells were added at clonal density by serial dilution from a 50,000 cells/ml stock. To account for variability in cell counting, dilution accuracy, and survival, a range of 20, 50, or 100 RFP⁺ cells per well was plated in parallel. Only wells in which RFP⁺ clones were clearly identifiable and spatially isolated were selected for analysis. Plates were fixed on day 28 and screened using an Incucyte microscope (Sartorius) to identify wells containing RFP⁺ clones. Positive wells were subsequently processed for immunostaining to identify PAX2+ dI4s and LMX1B+ dI5s (see Immunostaining section). Whole wells were scanned on an OperaPhenix microscope (Revvity) at 10X magnification, and clones were automatically identified based on RFP nuclear segmentation and local cell density. Regions containing candidate clones were then re-imaged at 40X NA1.1 for high-resolution analysis. Only discrete clones (Fig. S6D-E) were subsequently analysed. The number of LMX1B⁺/RFP⁺ and PAX2⁺/RFP⁺ cells was quantified based on intensity (using Harmony 5.0 software; Revvity) and manually curated to ensure accuracy. The ratio of PAX2⁺ to total PAX2⁺ + LMX1B⁺ cells was calculated per clone.

### Data and availability

All sequencing data associated with this study will be deposited in GEO (data submission pending; deposition delayed due to temporary suspension of NCBI services following a U.S. government funding lapse). The code supporting this study is available at: https://github.com/giulia-boezio/LARRY_spinal_cord_embryo_2025. The pipeline for LARRY clonal assignment is available at https://github.com/FrancisCrickInstitute/LARRY.

## Acknowledgements

We are grateful to Ariel Levine, Samantha Butler, Lora Sweeney, Joaquina Delás, Matteo Perino, and members of the Briscoe lab for their constructive feedback on the manuscript and Rubén Peréz- Carrasco for invaluable discussions. We thank Bertie Göttgens and Shirom Chabra for their input and troubleshooting support on the LARRY bioinformatics pipeline, and Emily Calderbank and Myriam Haltalli for assistance with LARRY library amplification protocols. We are also grateful to Fay Cooper and Anestis Tsakiridis for generating and sharing the H9-RFP line, to Manuela Melchionda for help with human embryo work, to Despina Stamataki and Ashley Libby for their help in establishing chicken protocols and in embryo dissection. We thank Fernando Camargo for providing the LARRY plasmid library. We acknowledge the following Science Technology Platforms at the Francis Crick Institute for their invaluable support: Genomics STP, Flow Cytometry STP, Advanced Light Microscopy STP, Vector Core, High Throughput Screening STP, and the Biological Research Facility.

## Author contributions

G.L.M.B. and J.B. conceived the project, interpreted data, and wrote the manuscript. G.L.M.B. designed and performed experiments. T.F. contributed to the design of *in vitro* hESCs differentiations and performed related experiments. J.D. and S.S. conceived the LARRY analysis pipeline. G.L.M.B., J.D., and A.R. performed bioinformatic analyses. M.H. contributed to the design of the *in vitro* clonal analyses. A.C. produced and validated the LARRY lentivirus.

---

**Figure S1.**
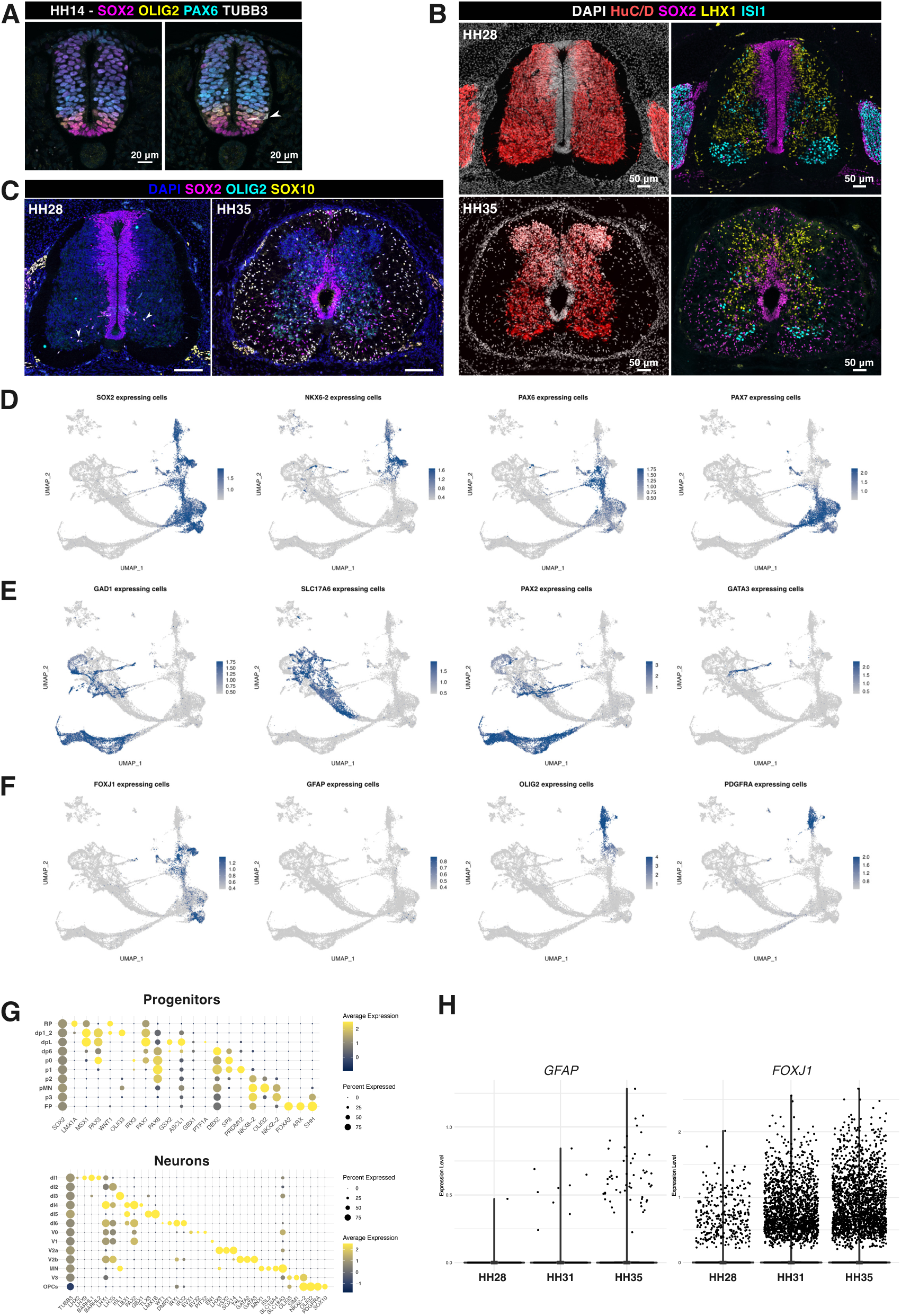
**Single-cell atlas of chicken spinal cord development** A. Confocal images of HH14 transverse sections stained for SOX2, OLIG2, PAX6, and TUBB3, showing progenitor patterning and near absence of TUBB3^+^ neurons. B. Confocal images of HH28 and HH35 transverse sections stained for HuC/D (pan-neural), SOX2 (neural and glial progenitors), LHX1, and ISL1, identifying diverse neuronal populations. C. Confocal image of HH28 and HH35 embryos transverse sections stained for SOX2, OLIG2, and SOX10. Triple- positive cells (arrowheads) in HH28 and HH35 sections indicate the emergence and expansion of OPCs. D–F. FeaturePlots showing marker expression for progenitors (D), neurons (E), and glia (F). G. Bubble plots showing expression of dorsoventral progenitor domain markers (top) and neuronal class markers (bottom); dot size indicates percentage of expressing cells, colour denotes average expression level. H. Boxplots of *FOXJ1* (ependymal cells) and *GFAP (*astrocytes) expression across developmental stages, showing progressive increase in expressing cells.

**Figure S2.**
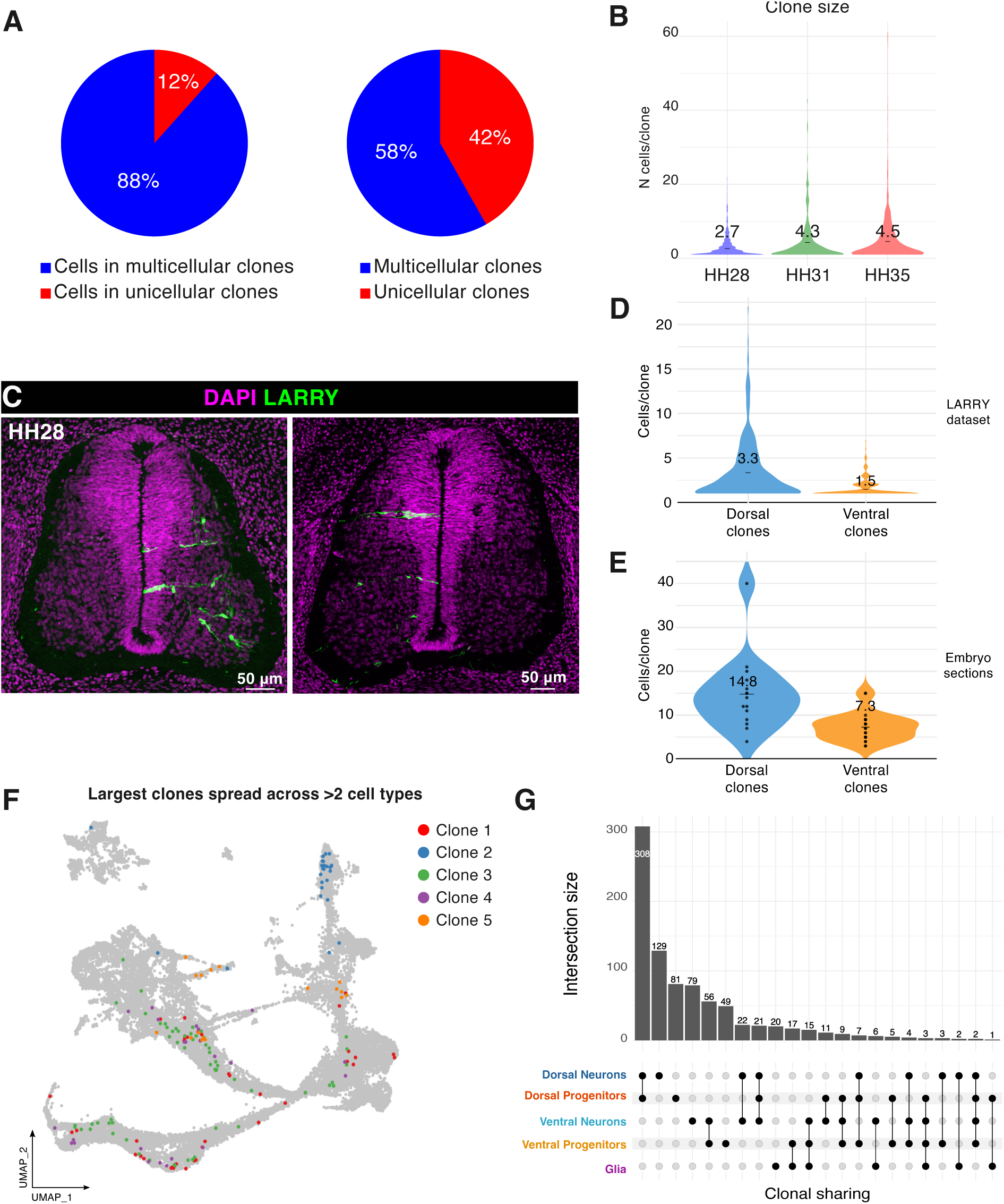
**LARRY barcoding in the chick spinal cord** A. Pie-charts showing percentage of cells in multicellular clones (left) and percentage of multicellular/single-cell clones (right). B. Violin plot of clone size distributions across datasets, highlighting the mean value. C. Confocal images of transverse spinal cord sections stained for EGFP (LARRY, green) showing individual ventral (left) and dorsal (right) clones in HH28 embryos. D-E. Violin plots of clone sizes in ventral and dorsal clones from scRNA-seq (D) and intact-tissue sections imaging (E), highlighting the mean. F. UMAP of the five largest clones spanning multiple cell types. G. UpSet plot displaying co-occurrence of cell types within individual clones. Top: bar plot quantifying clone combinations; bottom: graphical table showing the specific cell type combinations.

**Figure S3.**
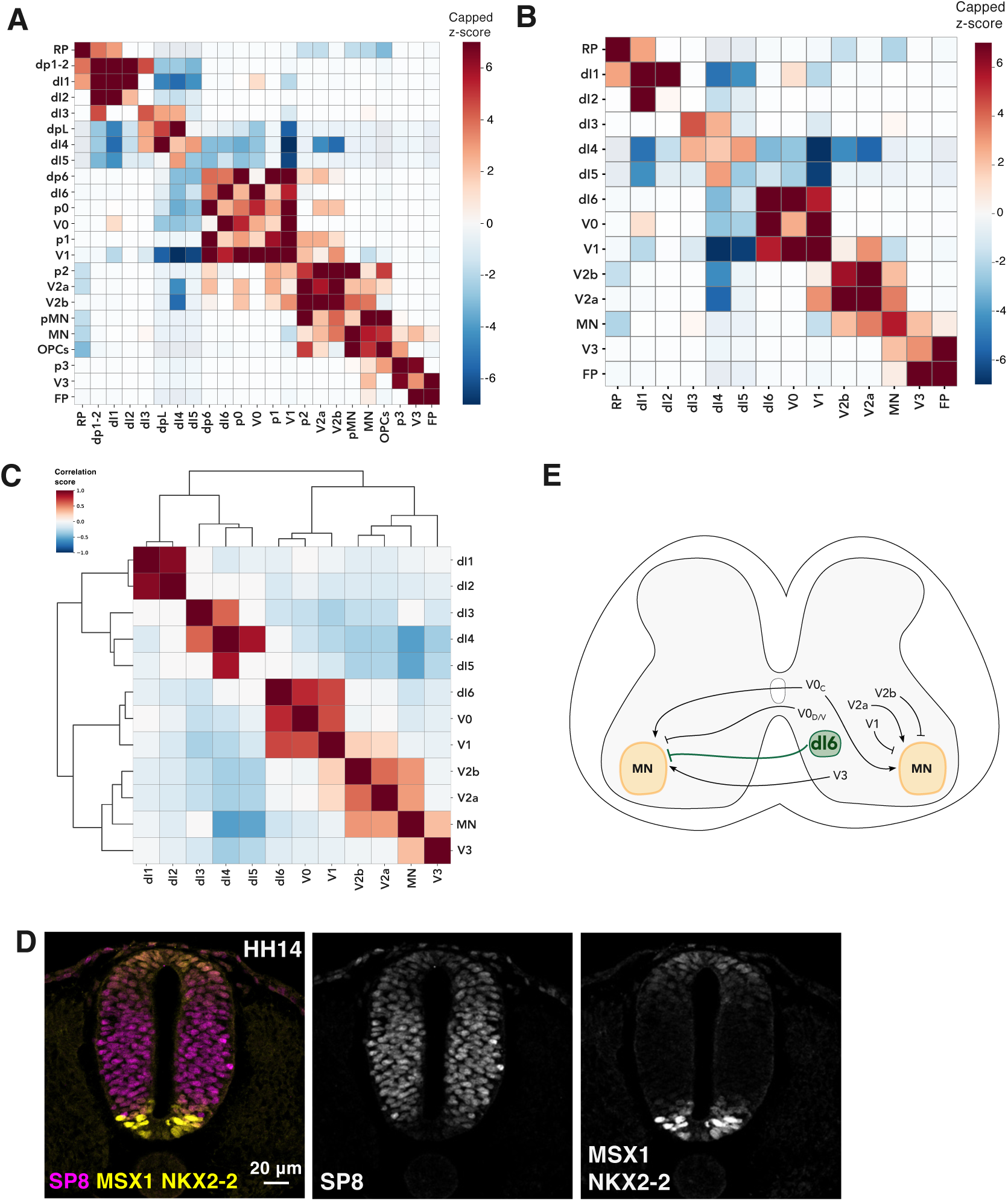
**Lineage tracing reveals five spatial lineage subdivisions** A-B. Heatmaps of clonal coupling z-scores for all cell types (A) and neurons only (B). Diagonal values indicate how frequently cells of the same type are found in the same clone; off-diagonal blocks highlight shared progenitor relationships between different cell types. C. Confocal images of HH14 transverse sections stained for different spatial transcription factors, corresponding to subdivisions in Fig. 2B. E. Diagram of functional connections between ventral interneurons and MNs, with emphasis on motor control roles of dI6 neurons.

**Figure S4.**
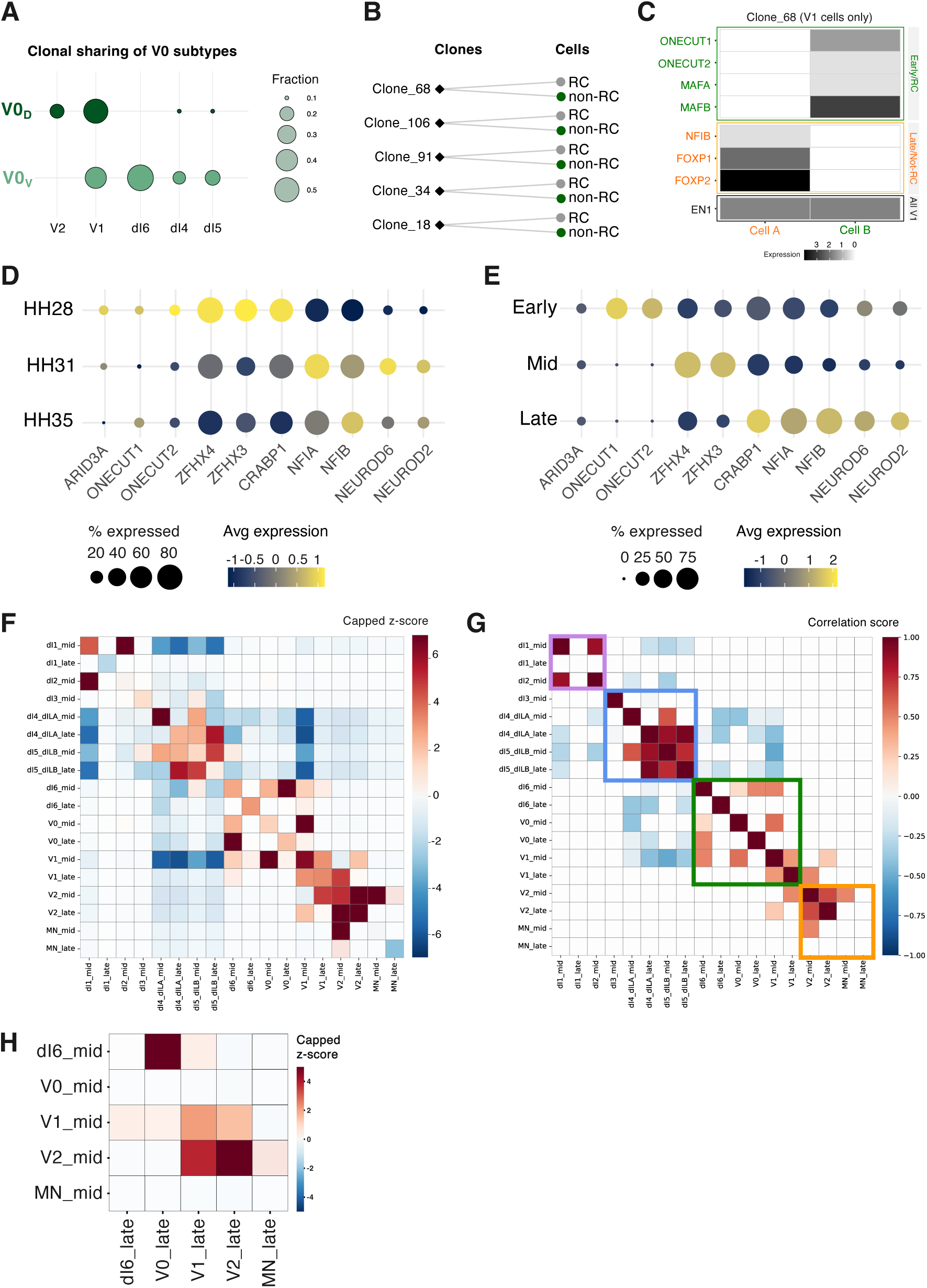
**Temporal dynamics of neurogenesis and clonal composition** A. Bubble plot showing the fraction of V0V and V0D clones that also contain other cell types. The radius of each bubble represents the fraction of total V0V and V0D clones. B. Bipartite plots show multicellular V1 clones (diamonds) linked to their constituent Renshaw (RC) and non- Renshaw (non-RC) neurons. C. Example V1 clone (Clone_68) with heatmaps of marker expression illustrating early RC identity (*ONECUT1/2, MAFB)* and late non-RC identity (*FOXP1/2, NFIB*). D-E. Bubble plots showing expression of temporal transcription factor markers across developmental datasets (D) and three neurogenic waves (E). Bubble colour represents the scaled average expression (z-score) of each transcription factor within the group. Values > 0 indicate above-average expression relative to all cells, and values < 0 indicate below-average expression. F-G. Heatmaps of clonal coupling z-scores (F) and correlation scores (G) among neurons stratified by temporal wave. Subdivisions are highlighted in G. H. Clonal coupling z-score heatmap for ventral neural subtypes, showing temporal progression from mid- to late-born neurons with a ventralisation trend.

**Figure S5.**
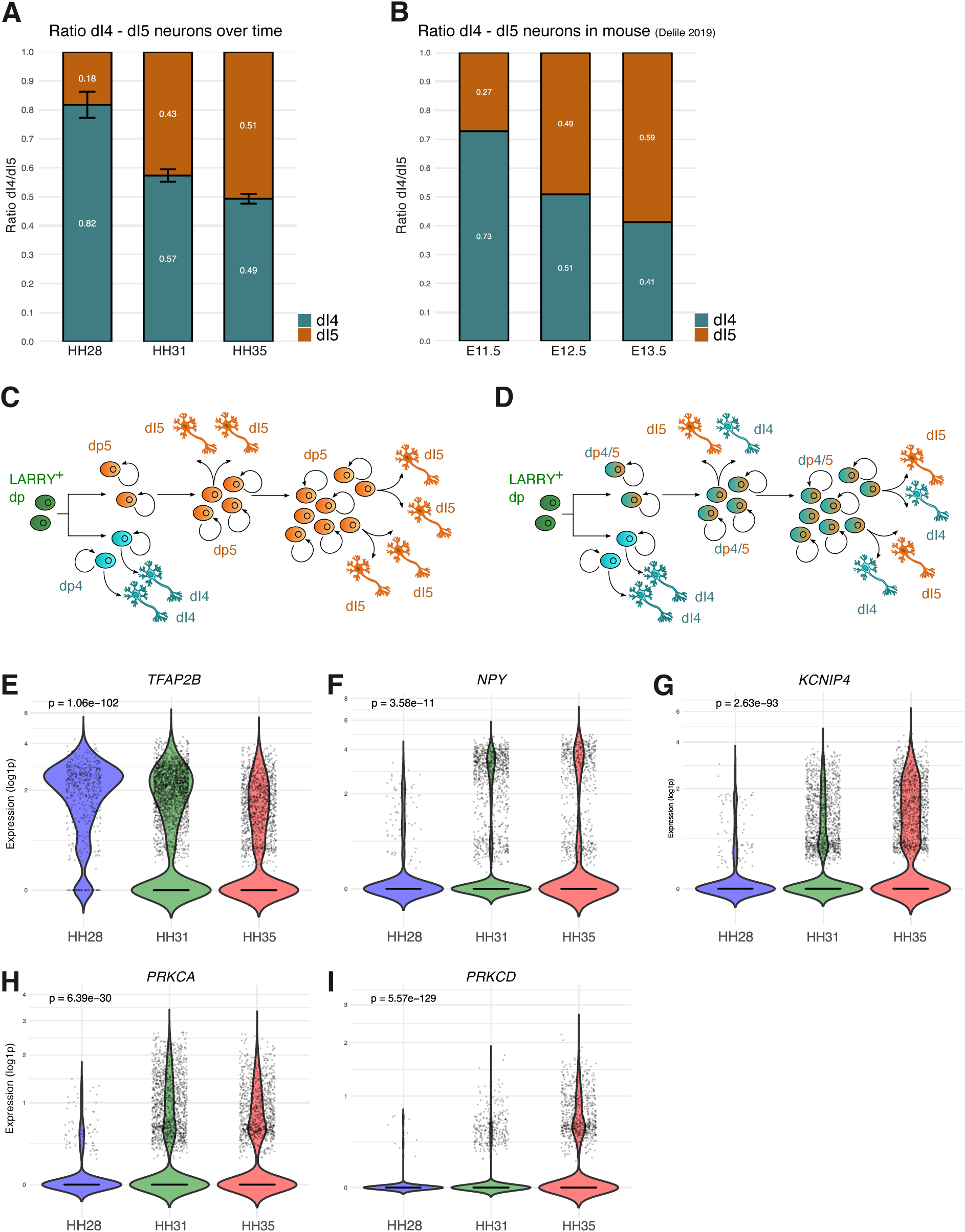
**Molecular characterisation and clonal dynamics of dI4/dI5 neurons** A-B. dI4/dI5 neuron ratios across timepoints in all cells (A) and in mouse neural tube datasets (B; *27*). C-D. Diagrams of potential lineage trajectories for dI4 and dI5 neurons. E-I. Violin plots showing expression levels and proportion of expressing cells for T*FAP2B, NPY*, *KCNIP4, PRKCA/D* over time.

**Figure S6.**
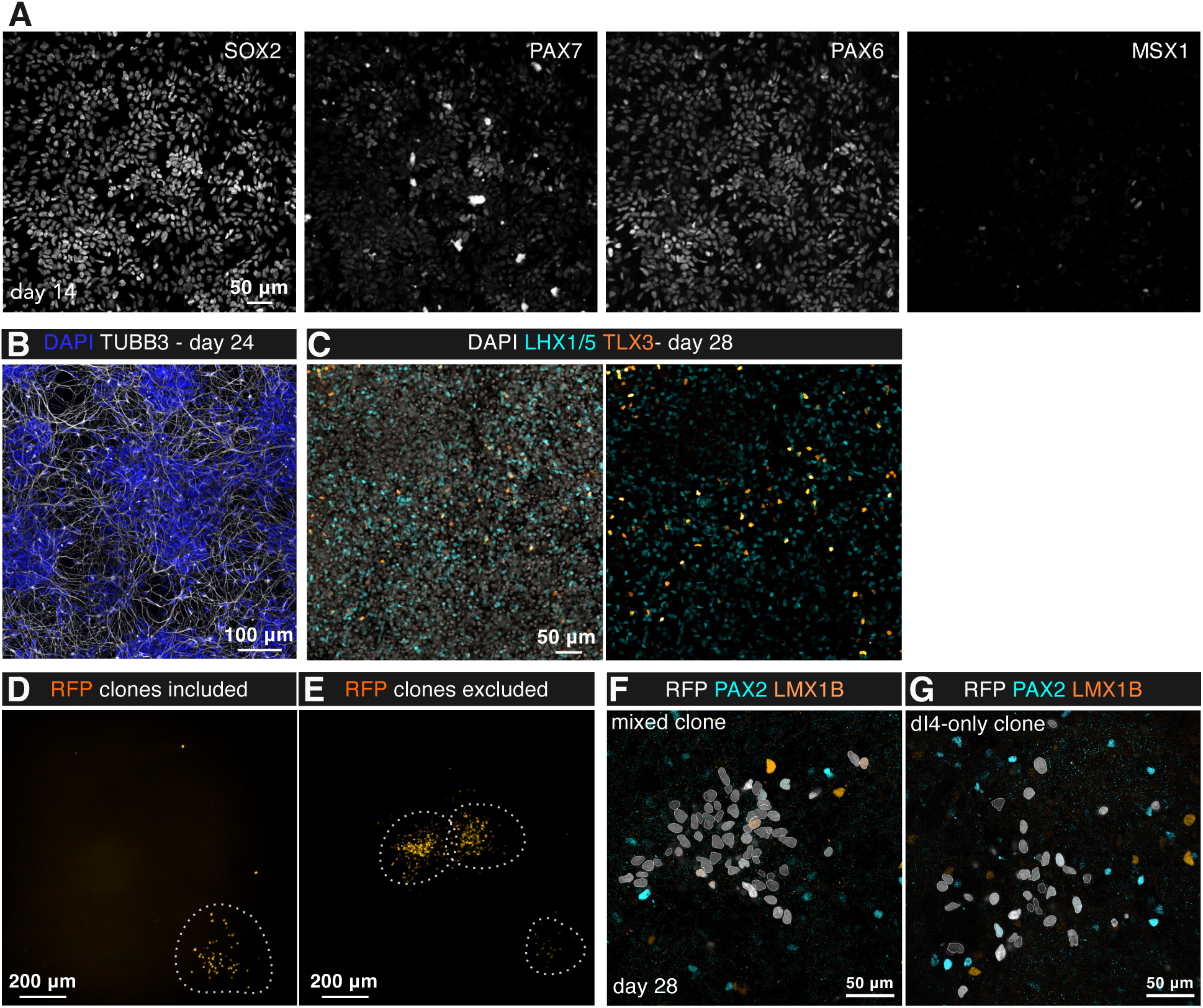
Characterisation of dI4/dI5 differentiation and clonal analysis in human *in vitro* models A. Confocal images of hESCs at day 14 showing dp4/dp5 progenitor identity (SOX2, PAX6, PAX7 positive). Cells are stained for SOX2, PAX6, PAX7, and MSX1. B. Confocal images of day 24 cultures stained for TUBB3 (βIII-tubulin, grey) showing extensive axonal outgrowth. C. Confocal images of day 28 cultures stained for LHX1/5 (cyan, dI4) and TLX3 (orange, dI5), confirming subtype-specific marker expression. D-E. Example of RFP-labelled clones at day 28. D; dotted lines outline confidently assigned clone used for quantification; E, dotted lines outline ambiguous RFP+ clusters excluded from analysis due to unclear clonal boundaries. F-G. Confocal images of day 28 H9-RFP clones showing examples of mixed clones (F) and dI4-only clones (G), relative to Figure 5E-F.

**Figure S7.**
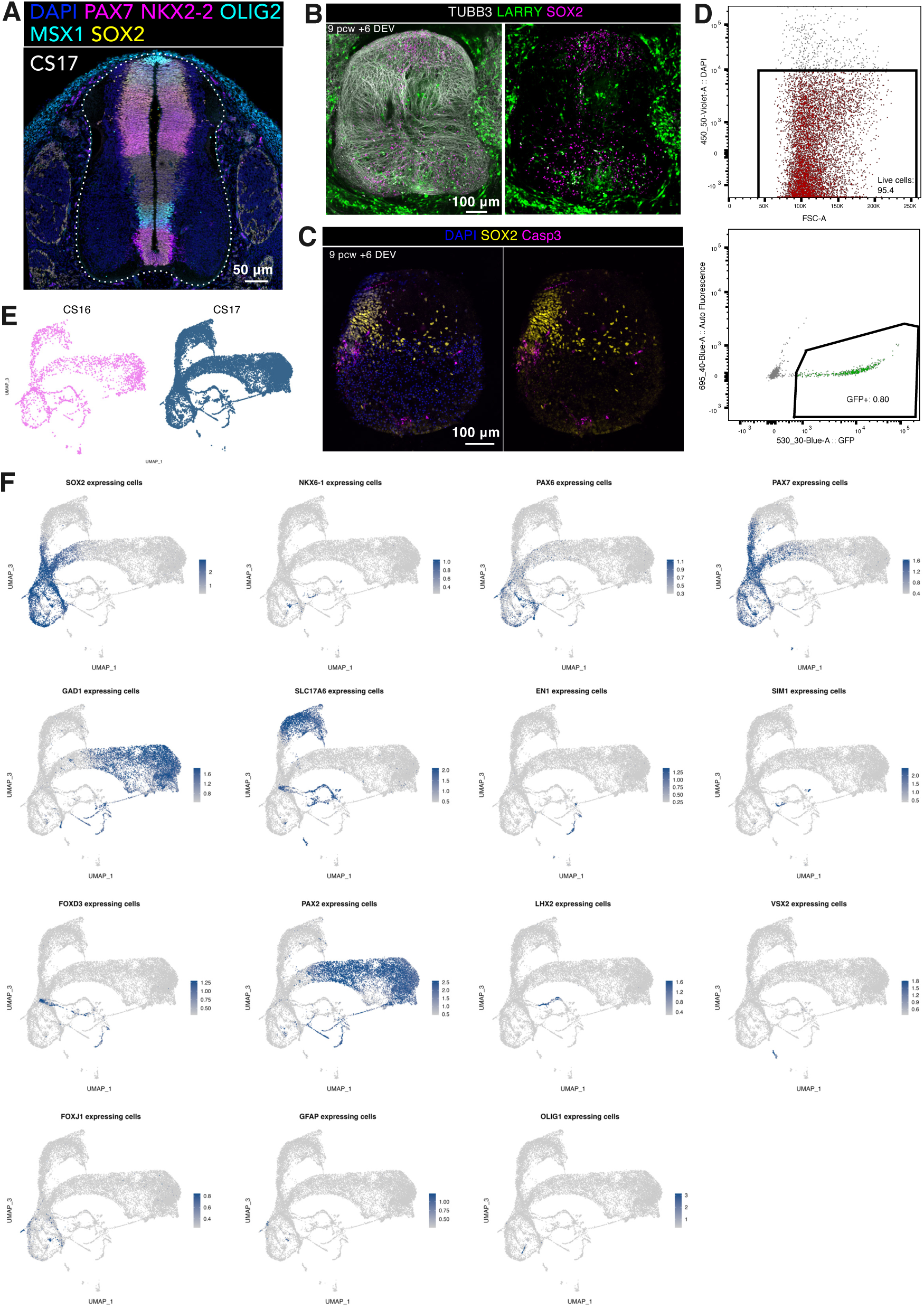
Validation of human spinal cord *ex vivo* cultures A. Confocal image of CS17 human spinal cord transverse section showing patterned progenitors in the ventricular zone. The shape of the spinal cord highlights a higher density of differentiated neurons (SOX2^-^) in the ventral regions, consistent with the earlier onset of ventral neurogenesis. B. Maximum intensity projections of 9 pcw + 6-day ex vivo (DEV) slices showing LARRY+ (green) progenitors (SOX2+, magenta) and neurons (TUBB3+, grey). C. Confocal images of 9 pcw + 6-day ex vivo (DEV) sections stained for Casp3 and SOX2, showing limited apoptosis in the tissue. D. Flow cytometry plots of CS17 + 7 DEV sample showing 95.4% viability and 0.8% GFP+ cells. E. UMAP of CS16 and CS17 samples. F. FeaturePlots of progenitor, neuron, and glia markers.

**Figure S8.**
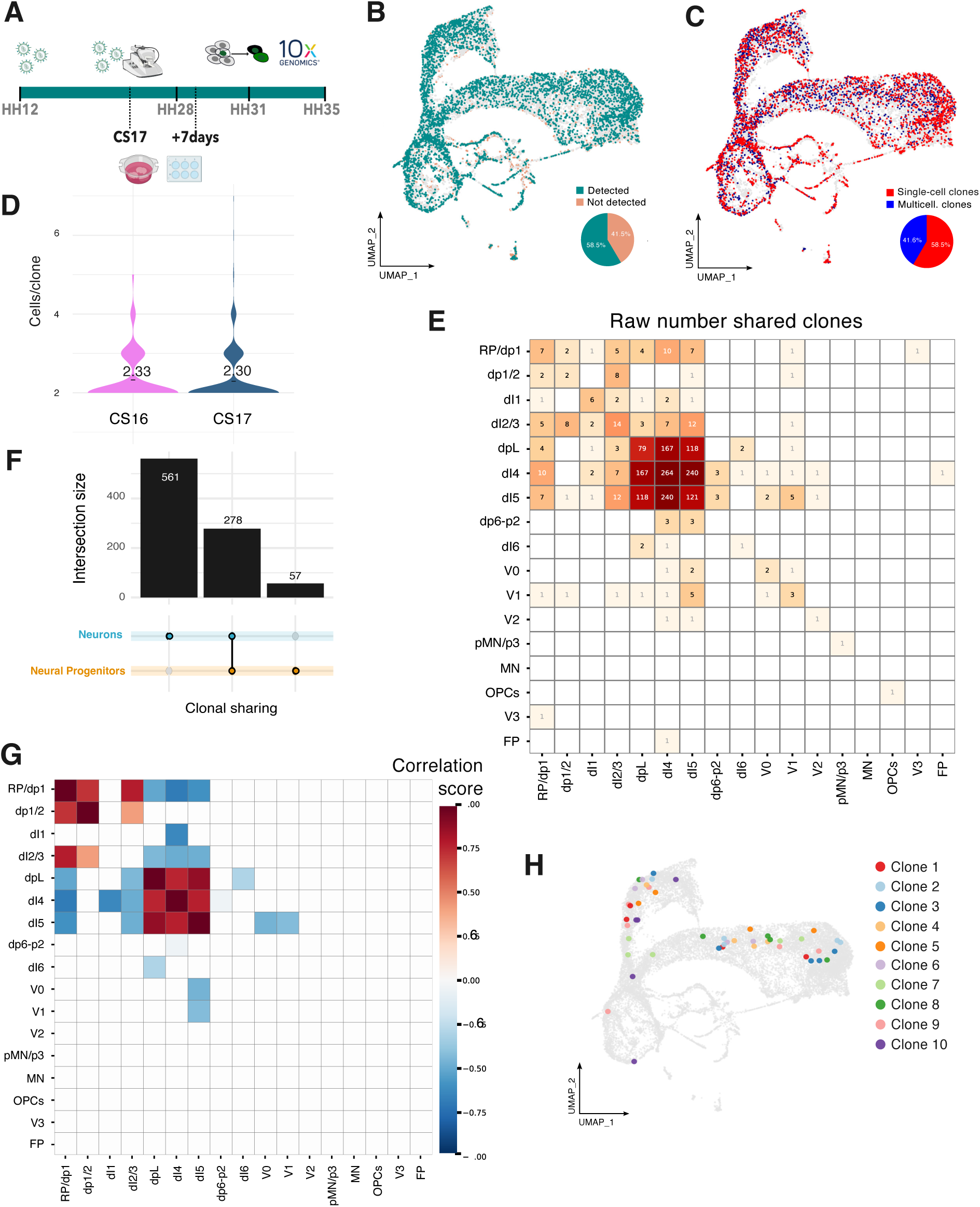
**Clonal lineage analysis in the human spinal cord** A. Comparative timeline of human and chicken stages of infection and collection. B-C. UMAPs of human samples showing barcode-expressing cells (B) and cells in multicellular clones (C). D. Violin plots of clone sizes across human samples. E. Heatmap of raw clone counts shared between cell type pairs. F. UpSet plot showing co-occurrence of cell types within clones. G. Heatmaps of clonal coupling correlation scores across all cell type pairs. H. UMAP of integrated human datasets showing the ten largest clones containing both dI4 and dI5 neurons.

**Figure S9.**
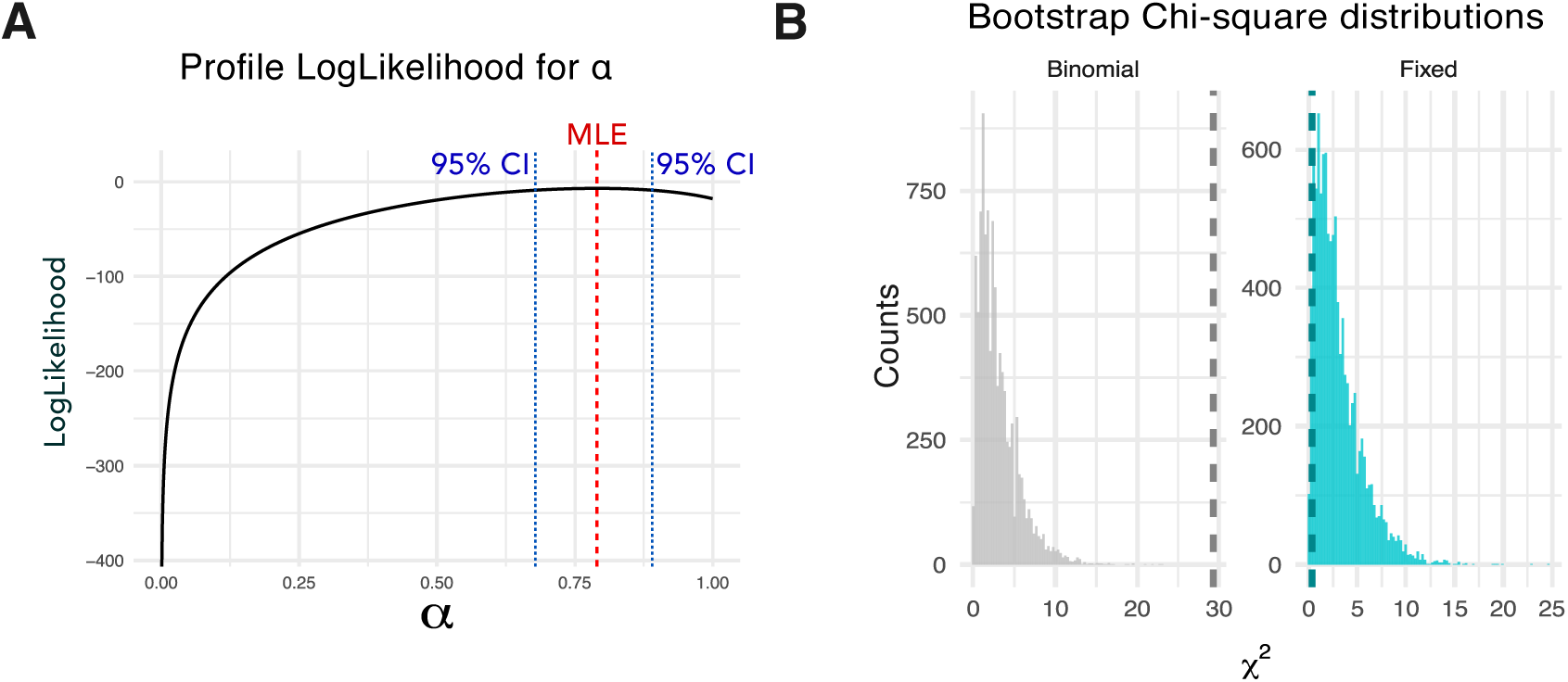
**dI4-dI5 lineage trajectories in the human embryonic spinal cord** A. Profile log-likelihood plot for the mixture weight α in the fixed mixture model. The maximum likelihood estimate (MLE, red) and 95% confidence interval (blue) are indicated. B. Distributions of χ² values from 10,000 bootstrapped datasets simulated under each model. The dashed line marks the χ² value from the actual data.

Table 1: Empirical p-values from bootstrapped χ² tests comparing observed and model-predicted clone distributions.

Table 2: Comparison of observed and model-predicted clone counts per bin, including bootstrapped 95% confidence intervals and observed–expected differences for each model.

Table 3: Model comparison metrics (log-likelihood, AIC, BIC) for binomial and fixed mixture models.

Table 4: List of molecular markers used for cell type annotation

Table 5: List of primers and sequencing indexes for LARRY library amplification from cDNA

